# DeepEthogram: a machine learning pipeline for supervised behavior classification from raw pixels

**DOI:** 10.1101/2020.09.24.312504

**Authors:** James P. Bohnslav, Nivanthika K. Wimalasena, Kelsey J. Clausing, David Yarmolinksy, Tomás Cruz, Eugenia Chiappe, Lauren L. Orefice, Clifford J. Woolf, Christopher D. Harvey

## Abstract

Researchers commonly acquire videos of animal behavior and quantify the prevalence of behaviors of interest to study nervous system function, the effects of gene mutations, and the efficacy of pharmacological therapies. This analysis is typically performed manually and is therefore immensely time consuming, often limited to a small number of behaviors, and variable across researchers. Here, we created DeepEthogram: software that takes raw pixel values of videos as input and uses machine learning to output an ethogram, the set of user-defined behaviors of interest present in each frame of a video. We used convolutional neural network models that compute motion in a video, extract features from motion and single frames, and classify these features into behaviors. These models classified behaviors with greater than 90% accuracy on single frames in videos of flies and mice, matching expert-level human performance. The models accurately predicted even extremely rare behaviors, required little training data, and generalized to new videos and subjects. DeepEthogram runs rapidly on common scientific computer hardware and has a graphical user interface that does not require programming by the end-user. We anticipate DeepEthogram will enable the rapid, automated, and reproducible assignment of behavior labels to every frame of a video, thus accelerating all those studies that quantify behaviors of interest.

Code is available at: https://github.com/jbohnslav/deepethogram

## Introduction

The analysis of animal behavior is a common approach in a wide range of biomedical research fields, including basic neuroscience research^1^, translational analysis of disease models, and development of therapeutics. For example, researchers study behavioral patterns of animals to investigate the effect of a gene mutation, understand the efficacy of potential pharmacological therapies, or uncover the neural underpinnings of behavior. In some cases, behavioral tests allow quantification of behavior through tracking an animal’s location in space, such as in the three-chamber assay, open field arena, Morris water maze, and elevated plus maze^2^. Increasingly researchers are finding that important details of behavior involve subtle actions that are hard to quantify, such as changes in the prevalence of grooming in models of anxiety^3^, licking a limb in models of pain^4^, and manipulation of food objects for fine sensorimotor control^5,6^. In these cases, researchers often closely observe videos of animals and then develop a list of behaviors they want to measure. To quantify these observations, by far the most commonly used approach, to our knowledge, is for researchers to manually watch videos with a stopwatch to count the time spent exhibiting each behavior of interest (Fig. 1A). This approach takes immense amounts of researcher time, often equal to or greater than the duration of the video per individual subject. Also, because this approach requires manual viewing, often only one or a small number of behaviors can be studied at a time. In addition, researchers often do not label the video frames when specific behaviors occur, precluding subsequent analysis and review of behavior bouts, such as bout durations and the transition probability between behaviors. Furthermore, scoring of behaviors can vary greatly between researchers especially as new researchers are trained^7^ and can be subject to bias. Therefore, it would be a significant advance if a researcher could define a list of behaviors of interest, such as face grooming, tail grooming, limb licking, locomoting, rearing, and so on, and then use automated software to identify when and how frequently each of the behaviors occurred in a video.

**Figure 1:**
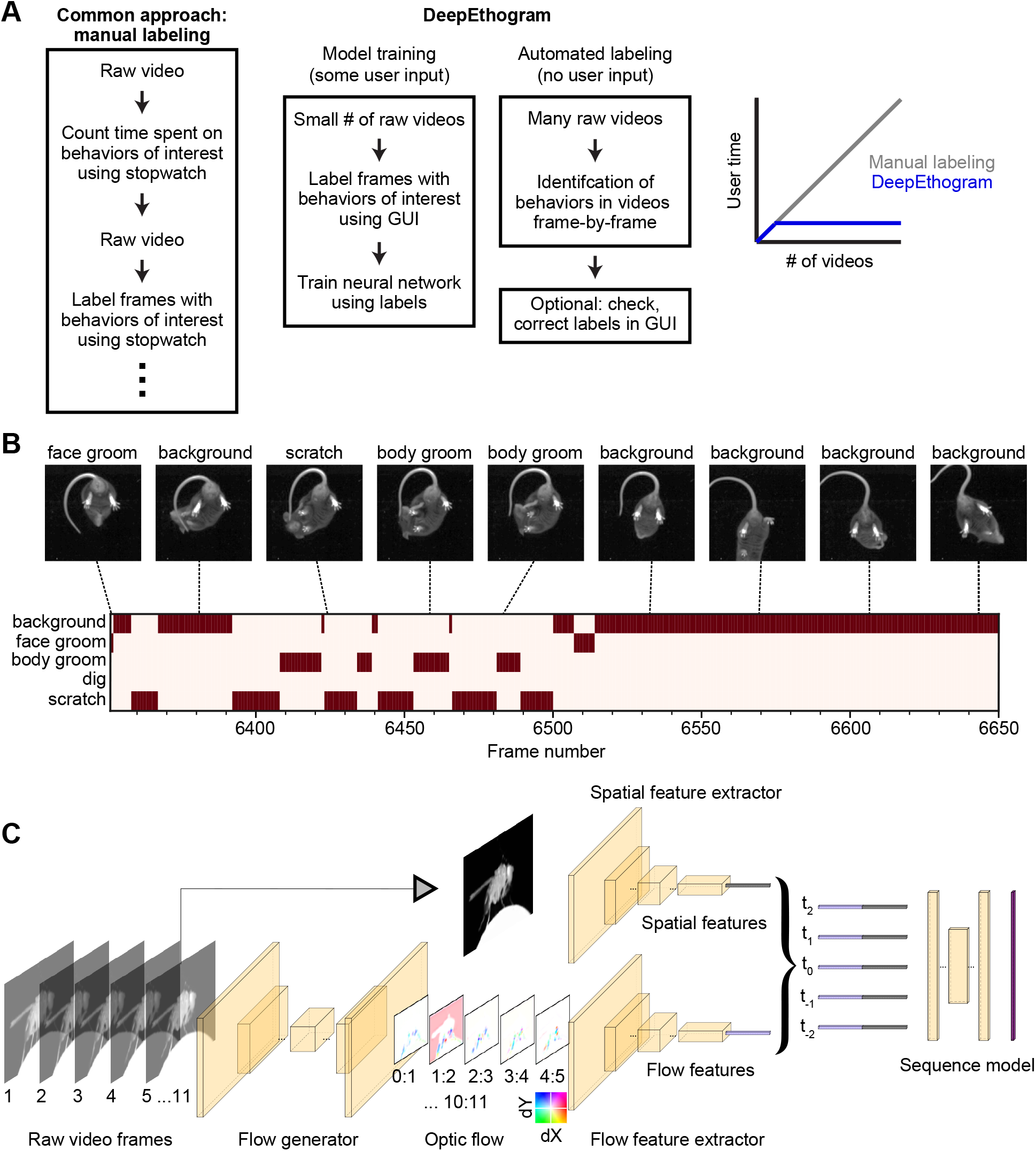
DeepEthogram overview. **(A)** Workflows for supervised behavior labeling. Left, a common traditional approach based on manual labeling. Middle, workflow with DeepEthogram. Right, Schematic of expected scaling of user-time for each workflow. **(B)** Ethogram schematic. Top, example images from Mouse-1 dataset. Bottom, ethogram with human labels. Dark colors indicate which behavior is present. Example shown is from Mouse-1 dataset. Images have been cropped, brightened, and converted to grayscale for clarity. **(C)** DEG-fast model schematic. Example images are from the Fly dataset. Left, a sequence of 11 frames is converted into 10 optic flows. Middle, the center frame and the stack of 10 optic flows are converted into 512-dimensional representations via deep CNNs. Right, these features are converted into probabilities of each behavior via the sequence model.

Researchers are increasingly turning to computational approaches to quantify and analyze animal behavior^8–12^. The task of automatically classifying an animal’s actions into user-defined behaviors falls in the category of supervised machine learning. In computer vision, this task is called “action detection”, “temporal action localization”, or “action segmentation”. This task is distinct from other emerging behavioral analysis methods based on unsupervised learning, in which machine learning models discover behavioral modules from the data, irrespective of researcher labels. Although unsupervised methods, such as Motion Sequencing^13,14^, MotionMapper^15^, BehaveNet^16^, B-SOiD^17^, and others^8^, can discover behavioral modules not obvious to the researcher, their outputs can be challenging to match up to behaviors of interest in cases in which researchers have strong prior knowledge about the specific behaviors relevant to their experiments.

Pioneering work, including JAABA^18^, SimBA^19^, MARS^7^, and others^7,20,21^, has made important progress toward the goal of supervised classification of behaviors. These methods track specific features of an animal’s body and use the time series of these features to classify whether a behavior is present at a given timepoint. In computer vision, this is known as “skeleton-based action detection”. In JAABA, ellipses are fit to the outline of an animal’s body, and these ellipses are used to classify behaviors. SimBA classifies behaviors based on the positions of “keypoints” on the animal’s body, such as limb joints, which are derived from other processing pipelines, such as DeepLabCut^22,23^ or similar algorithms^24,25^. MARS takes a similar approach, with a focus on social behaviors^7^. Thus, with these approaches, researchers first reduce a video to a set of user-defined features of interest and then classify behaviors based on those features. This approach has four important limitations. First, the user must specify which features are key to the behavior (e.g. body position or limb position), but many behaviors are whole-body activities that could best be classified by full body data. Second, errors that occur in tracking these features in a video will result in poor input data to the classification of behaviors, potentially decreasing the accuracy of labeling. Third, users might have to perform a pre-processing step between their raw videos and the input to these algorithms, increasing pipeline complexity and researcher time. Fourth, the selection of features often needs to be tailored to specific video angles, behaviors (e.g. social behaviors vs. individual mice), species, and maze environments, making the analysis pipelines often specialized to specific experiments. Other recent work uses image and motion features, similar to the approaches we developed here, except with a focus on classifying the timepoint at which a behavior starts, instead of classifying every frame into one or more behaviors^26^. Still other supervised learning algorithms, such as DeepLabCut^22,23^, LEAP^27^, DeepPoseKit^25^, and others^8^ track specific body features (e.g. limb movements), without classifying these movements into behaviors, such as face grooming, body grooming, or limb licking, and thus are not the same as automated behavior classification.

Here we developed a modular pipeline for automatically classifying each frame of a video into a set of user-defined behaviors. Our method, called DeepEthogram (DEG), uses a supervised deep-learning model that, with minimal user-based training data, takes a video with *T* frames and a user-defined set of *K* behaviors and generates a binary [*T, K*] matrix (Fig. 1A). This matrix indicates whether each behavior is present or absent at each frame, which we term an “ethogram”: the set of behaviors that are present at a given timepoint (Fig. 1B). Our model operates directly on the raw pixel values of videos, and thus it is generally applicable to any case with video data and binary behavior labels and further does not require pre-specification of the body features of interest, such as keypoints on limbs or fitting the body with ellipses. We used state-of-the-art convolutional neural networks, specifically Hidden Two-Stream Networks^28^ and Temporal Gaussian Mixture networks^29^, to detect actions in videos and pre-trained the networks on large open-source datasets^30,31^. We validated our approach’s performance on four datasets from two species, with each dataset posing distinct challenges for behavior classification. DEG automatically classified behaviors with high performance, often reaching levels obtained by expert human labelers. High performance was achieved with only a few minutes of positive example data and even when the behaviors occurred at different locations in the behavioral arena and at distinct orientations of the animal. Importantly, the entire pipeline requires no programming by the end-user because we developed a graphical user-interface (GUI) for annotating videos, training models, and generating predictions.

## Results

### Modeling approach

Our goal was to take a set of video frames as input and predict the probability that each behavior of interest occurs on a given frame. This task of automated behavior labeling presents several challenges that framed our solution. First, in many cases, the behavior of interest occurs in a relatively small number of video frames, and the accuracy must be judged based on correct identification of these low frequency events. For example, if a behavior of interest is present in 5% of frames, an algorithm could guess that the behavior is “not present” on every frame and still achieve 95% overall accuracy. Critically, however, it would achieve 0% accuracy on the frames that matter, and an algorithm does not know a priori which frames matter. Second, ideally a method should perform well after being trained on only small amounts of user-labeled video frames, including across different animals, and thus require little manual input. Third, a method should be able to identify the same behavior regardless of the position and orientation of the animal when the behavior occurs. Fourth, methods should require relatively low computational resources in case researchers do not have access to large compute clusters or top-level GPUs.

We modeled our approach after temporal action localization methods used in computer vision aimed to solve related problems^32–35^. The overall architecture of our solution included: 1. estimating motion (optic flow) from a small snippet of video frames, 2. compressing a snippet of optic flow and individual still images into a lower dimensional set of features, 3. using a sequence of the compressed features to estimate the probability of each behavior at each frame in a video (Fig. 1C). We implemented this architecture using large, very deep convolutional neural networks (CNNs). First, one CNN was used to generate optic flow from a set of images. We incorporated optic flow because some behaviors are only obvious by looking at the animal’s movements between frames. For example, one can distinguish standing still and walking using a short snippet of the animal’s movements. We call this CNN the *flow generator* (Fig. 1C, Supplementary Fig. 1). We then used the optic flow output of the flow generator as input to a second CNN to compress the large number of optic flow snippets across all the pixels into a small set of features called *flow features* (Fig. 1C). Separately, we used a distinct CNN, which takes single video frames as input, to compress the large number of raw pixels into a small set of *spatial features*, which contain information about the values of pixels relative to one another spatially but lack temporal information (Fig. 1C). We included single frames separately because some behaviors are obvious from a single still image from a video, such as identifying licking just by seeing an extended tongue. Together, we call these latter two CNNs *feature extractors* because they compressed the large number of raw pixels into a small set of features called a *feature vector* (Fig. 1C). Each of these feature extractors was trained to produce a probability for each behavior on each frame based only on their input (optic flow or single frames). We then fused the outputs of the two feature extractors by averaging (Methods). To produce the final probabilities that each behavior was present on a given frame – a step called inference – we used a *sequence model*, which has a large temporal receptive field and thus utilizes long timescale information (Fig. 1C). We used this sequence model because our CNNs only “looked at” either one frame (spatial) or about 11 frames (optic flow), but when labeling videos, humans know that sometimes the information present seconds ago can be useful for estimating the behavior of the current frame. The final output of DeepEthogram (DEG) is a [*T, K*| matrix, in which each element is the probability of behavior *k* occurring at time *t*. We thresholded these probabilities to get a binary prediction for each behavior at each time point, with the possibility that multiple behaviors could occur simultaneously (Fig. 1B).

For the flow generator, we used the MotionNet^28^ architectures to generate ten optic flow frames from eleven images. For the feature extractors, we used the ResNet family of models^36,37^ to extract both flow features and spatial features. Finally, we used Temporal Gaussian Mixture^29^ models as the sequence model to perform the ultimate classification. Each of these models have many variants with a large range in the number of parameters and the associated computational demands. We therefore created three versions of DEG that use variants of these models and trade off accuracy and speed: DEG-fast, DEG-medium, and DEG-slow. DEG-fast uses TinyMotionNet^28^ for the flow generator and ResNet18^36^ for the feature extractors. It has the fewest parameters, the fastest training of the flow generator and feature extractor models, the fastest inference time, and the smallest requirement for computational resources. As a tradeoff for this speed, DEG-fast also has the worst performance (see below). In contrast, DEG-slow uses a novel architecture TinyMotionNet3D for its flow generator and 3D-ResNet34^28,36,37^ for its feature extractors. It has the most parameters, the slowest training and inference times, and the highest computational demands, but it produces the best performance. DEG-medium is intermediate between DEG-fast and DEG-slow and uses MotionNet^28^ and ResNet50^36^ for its flow generator and feature extractors. All versions of DEG use the same sequence model. All variants of the flow generators and feature extractors were pretrained on the Kinetics700 video dataset^31^, so that model parameters did not have to be learned from scratch (Methods).

In practice, the first step in running DEG is to train the flow generator on a set of videos, which occurs without user input (Fig. 1A). In parallel, a user must label each frame in a set of training videos for the presence of each behavior of interest. These labels are then used to train independently the spatial feature extractor and the flow feature extractor to produce separate estimates of the probability of each behavior. The extracted feature vectors for each frame are then saved and used to train the sequence models to produce the final predicted probability of each behavior at each frame. We chose to train the models in series, rather than all at once from end-to-end, due to a combination of concerns about backpropagating error across diverse models, overfitting with extremely large models, and computational capacity (Methods).

### Diverse and challenging datasets to test DeepEthogram

To test the performance of our model, we used four different neuroscience research datasets. The datasets were selected because they spanned two species, and each presented distinct challenges for computer vision approaches. We specifically did not choose videos that may allow easy automatic classification. Please see the examples in Figure 2, Supplementary Figures S2-S3, and Supplementary Videos 1-4 that highlight these challenges in distinguishing one behavior from another.

**Figure 2:**
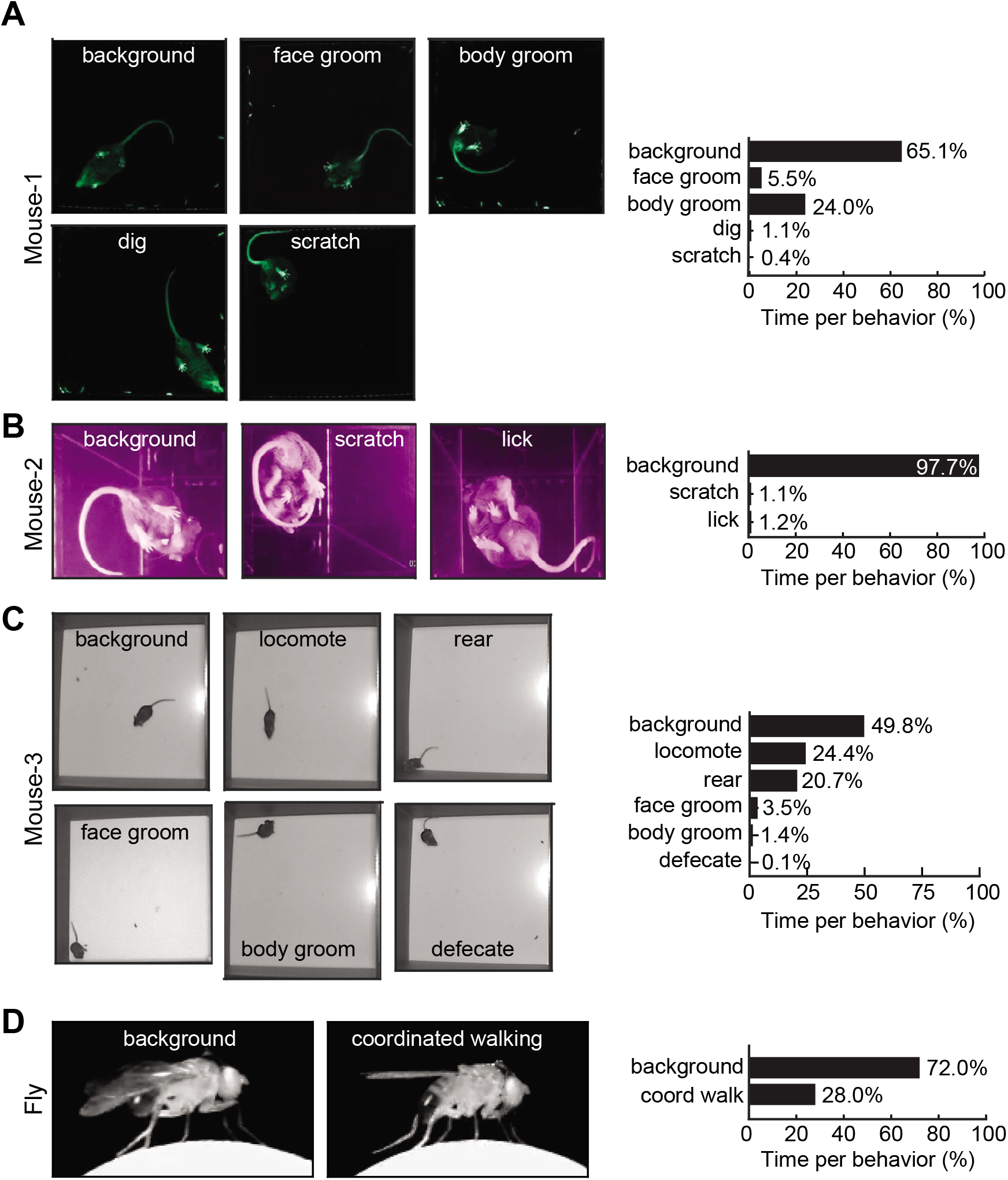
Datasets and behaviors of interest. **(A)** Left, raw example images from the Mouse-1 dataset for each of the behaviors of interest. Right, time spent on each behavior, based on human labels. Note that the times may add up to more than 100% across behaviors because multiple behaviors can occur on the same frame. Background is defined as the behavior when no other specified behaviors are occurring. **(B-D)** Similar to (A), except for the Mouse-2 dataset, Mouse-3 dataset, and Fly dataset.

The first dataset, called “Mouse-1”, is a bottom-up view of a mouse in a large open field box walking on a transparent platform (Fig. 2A, Supplementary Fig. 2A, Supplementary Video 1). We labeled each frame in these videos for the presence of four behaviors of interest (or classes): face grooming, body grooming, digging, and scratching. Following computer vision convention, we labeled the frames on which none of these behaviors were present as “background.” We chose these videos for the challenges that the mouse was generally poorly illuminated, and the mouse occupied a small number of pixels. An important feature of this dataset is that a given behavior, such as face grooming, could occur in any location in the open field and with the mouse facing in any direction. In all cases, we labeled these frames as examples of the “face grooming” class and did not note any position or direction information. As such, we did not spatially crop the video frames or align the animal before training our model.

The second dataset, called “Mouse-2”, is another bottom-up view of a mouse, except in a small chamber (Fig. 2B, Supplementary Fig. 2B, Supplementary Video 2). The behaviors of interest were scratch and lick. These videos were chosen because they present the challenge of being relatively low quality due to compression artifacts, reflections of the mouse on the walls of the chamber, and large distortions from the camera lens. Also, the behaviors of interest occurred extremely infrequently, in only about 2% of the total video frames. This dataset therefore tests the model’s ability to successfully distinguish rare behaviors with non-ideal video quality.

The third dataset, called “Mouse-3”, is a top-down view of a mouse in an open chamber (Fig. 2C, Supplementary Fig. 3A, Supplementary Video 3). We chose these videos because they are typical in the analysis of mouse behavior, such as for the commonly used open field assay and other assays in which mice explore open environments. The behaviors of interest in this dataset were face grooming, body grooming, locomoting, rearing, and defecating. These videos presented the challenge of poor contrast between parts of the mouse’s body, making it difficult to determine the location of body parts. Also, the mouse occupied few pixels in the video, and the videos were relatively low resolution. This dataset therefore tests the model’s ability on typical mouse behavior analysis videos.

The fourth dataset, called “Fly”, is a side-view of a *Drosophila melanogaster* walking on a ball (Fig. 2D, Supplementary Fig. 3B, Supplementary Video 4). Previous work has demonstrated that neural activity is modulated when the fly exhibits stereotypical, coordinated walking^38^. Therefore, this dataset consists of one behavior of interest, coordinated walking. This dataset presents the challenge of having little motion information because the videos were recorded at a high frame rate, resulting in little frame-to-frame differences. This dataset thus tests our model’s ability to integrate information over time.

### DeepEthogram achieved high performance approaching expert-level human performance

We split each dataset into three subsets: training, validation, and test (Methods). The training set was used to update model parameters, such as the weights of the CNNs. The validation set was used to set appropriate hyperparameters, such as the thresholds used to turn the probabilities of each behavior into binary predictions about whether each behavior was present. The test set was used to report performance on new data not used in training the model. We computed three complementary metrics of model performance using the test set. First, we computed the accuracy, which is the fraction of elements of the [*T, K*| ethogram that were predicted correctly. We note, however, that in theory accuracy could be high even if the model did not perform well on each behavior. For example, in the Mouse-2 dataset, some behaviors were incredibly rare, occurring in only ~2% of frames (Fig. 2B). Thus, the model could in theory achieve ~98% accuracy simply by guessing that the behavior was absent on all frames. Therefore, we also computed the F1 score, a metric ranging from 0 (bad) to 1 (perfect) that takes into account the rates of false positives and false negatives. The F1 scores is the geometric mean of the precision and recall of the model. Precision is the fraction of frames labeled by the model as a given behavior that are actually that behavior (true positives / (true positives + false positives)). Recall is the fraction of frames actually having a given behavior that are correctly labeled as that behavior by the model (true positives / (true positives + false negatives)). We report the F1 score in the main figures and show precision and recall performance separately in the supplementary figures. Because the accuracy and F1 score depended on our choice of a threshold to turn the probability of a given behavior on a given frame into a binary prediction about the presence of that behavior, we also computed the area under the receiver operating curve (AUROC), which summarizes performance as a function of the threshold.

We first considered the entire ethogram, including all behaviors. DEG performed with greater than 90% accuracy on the test data for all datasets (Fig. 3A). Also, the model achieved very high overall F1 scores, with high precision and recall (Fig. 3B, Supplementary Figs. 4A, 5A). Similarly, high overall performance was observed with the AUROC measures (Supplementary Fig. 6A). These results indicate that the model was able to capture the overall patterns of behavior in videos.

**Figure 3:**
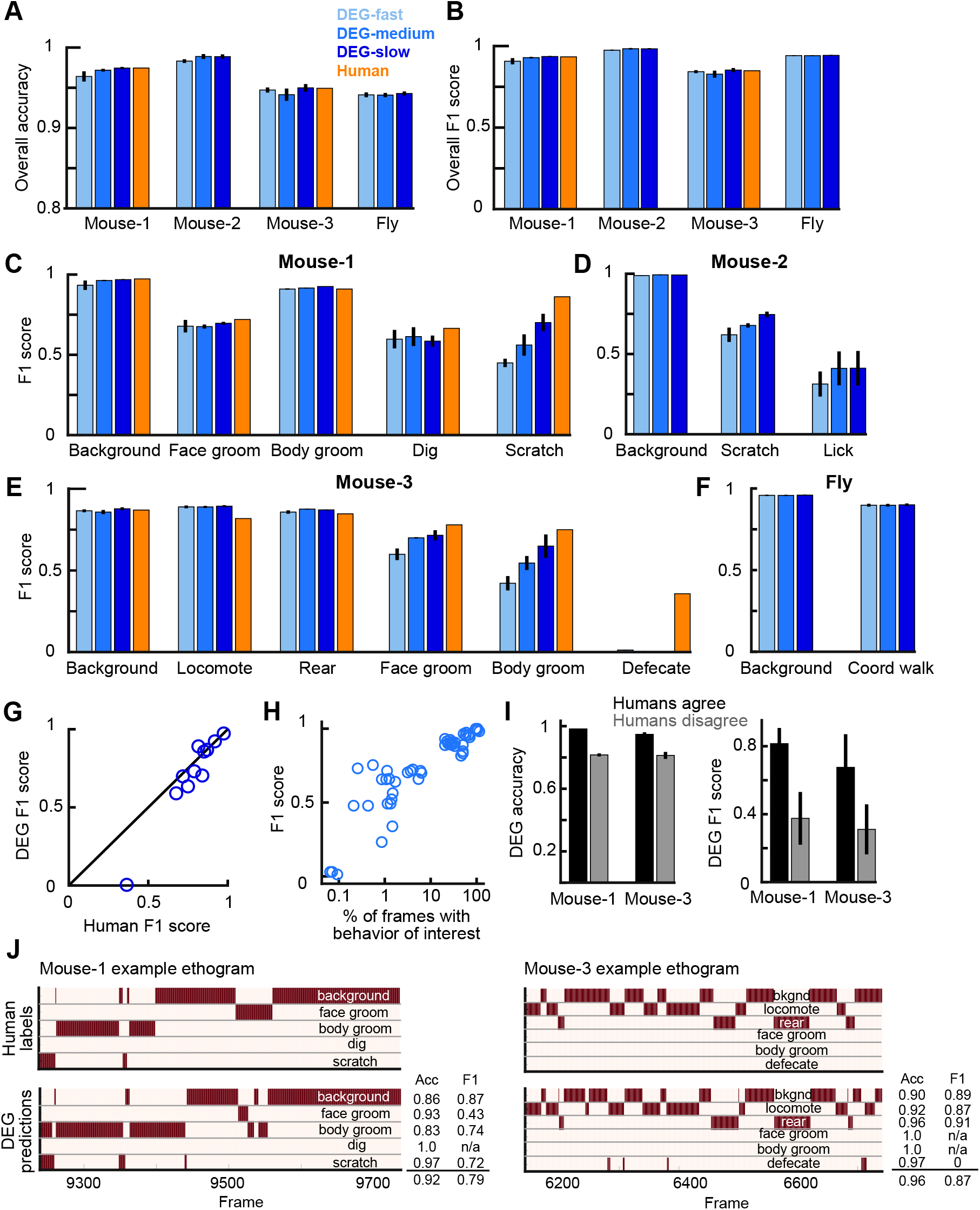
DEG performance. **(A)** Overall accuracy for each DEG variant and dataset for data from the test set. Human accuracy corresponds to the agreement between human labelers. Error bars indicate mean +/- s.e.m. across three random splits of the data. **(B)** Similar to (A), except for overall F1 score. **(C)** F1 score for each DEG variant for individual behaviors from the Mouse-1 dataset. **(D-F)** Similar to (C), except for the Mouse-2 dataset, Mouse-3 dataset, and Fly dataset. **(G)** F1 scores between two human labelers compared to DEG-slow. Each point is one behavior, with performance averaged across 3 random splits of the data. The point with low DEG F1 score is for “defecation” from Mouse-3. **(H)** DEG-medium performance measured as F1 score as a function of the percent of frames containing the behavior of interest in the training data. Each point is for one behavior for one split of the data, with three splits in total. **(I)** DEG-medium performance on frames when the two human labelers agreed or disagreed on the label. Error bars indicate mean +/- s.e.m. across three random splits of the data. **(J)** Ethogram examples. Dark color indicates labels for the behavior being present on a given frame. Top, human labels. Bottom, DEG-medium predictions. The accuracy and F1 score for each behavior, and the overall accuracy and F1 scores are shown. Examples were chosen to be similar to the model’s average by behavior (C-F).

We also analyzed the model’s performance for each individual behavior. The model achieved F1 scores of 0.7 or higher for many behaviors, even reaching F1 scores above 0.9 in some cases (Fig. 3C-F). Given that F1 scores may not be intuitive to understand in terms of their values, we examined individual snippets of videos with a range of F1 scores and found that F1 scores similar to the means for our datasets were consistent with overall accurate predictions (Fig. 3J, Supplementary Figs. 7-10). We note that the F1 score is a demanding metric, and even occasional differences on single frames or a small number of frames can substantially decrease the F1 score. Relatedly, the model achieved high precision, recall, and AUROC values for individual behaviors (Supplementary Figs. 4B-E, 5B-E, 6B-E). The performance of the model depended on the frequency with which a behavior occurred (c.f. Fig.2 (right panels) and Fig. 3C-F). Strikingly, however, performance was relatively high even for behaviors that occurred extremely rarely, i.e. in less than 10% of video frames (Fig. 3H, Supplementary Figs. 4G, 5G, 6F). The performance was generally highest for DEG-slow and worst for DEG-fast (Fig. 3C-F). This difference between DEG variants was less apparent on frequent behaviors and was largest on the rarest behaviors in the datasets. The high performance values are, in our opinion, especially impressive given that they were calculated based on single frame predictions for each behavior, and thus performance will be reduced even if the model misses the onset or offset of a behavior bout by even a single frame. These high values suggest that the model not only correctly predicted which behaviors happened and when, but also had the resolution to correctly predict the onset and offset of bouts.

To better understand the performance of DEG, we benchmarked the model by comparing its performance to the degree of agreement between expert human labelers. Two researchers with extensive experience in monitoring and analyzing mouse behavior videos independently labeled the same set of videos for the Mouse-1 dataset and a subset of the Mouse-3 dataset, allowing us to understand the consistency across human experts. Human-human performance was calculated by defining one labeler as the “ground truth” and the other labeler as “predictions”, and then computing the same performance metrics as for DEG. Strikingly, the overall accuracy, F1 scores, precision, and recall across these datasets were similar for DEG and human labelers (Fig. 3A,B,C,E,G, Supplementary Figs. 4A,B,D,F, 5A,B,D,F). DEG-slow approached human-level performance even on rare behaviors. DEG-fast and DEG-medium performed similarly to expert humans on most behaviors but performed worse on the rarest behaviors in each dataset. Notably, the behaviors for which DEG had the lowest performance tended to also be the behaviors for which humans had less agreement (lower human-human F1 score) (Fig. 3G, Supplementary Figs. 4F, 5F). Relatedly, DEG performed best on the frames in which the human labelers agreed and tended to do more poorly in the frames in which humans disagreed (Fig. 3I). Thus, there was a strong correlation between DEG and human performance, and the values for DEG performance were similar to those of expert human labelers, even for the rarest and most challenging behaviors (Fig. 3G).

The only behavior the model failed to identify was “defecate” from the Mouse-3 dataset (Figs. 2C, 3E). Notably, defecation was incredibly rare, occurring in only 0.1% of frames. Furthermore, the act of defecation was not actually visible from the videos. Rather, human labelers marked the “defecate” behavior when new fecal matter appeared, which involved knowledge of the foreground and background, tracking objects, and inferring unseen behavior. This type of behavior is expected to be challenging for DEG because DEG uses RGB images and local motion cues and thus will fail when the behavior cannot be directly observed visually.

The model was able to accurately predict the presence of a behavior even when that behavior happened in different locations in the environment and with different orientations of the animal (Supplementary Fig. 11). For example, the model predicted face grooming accurately both when the mouse was in the top left quadrant of the chamber and facing north and when the mouse was in the bottom right quadrant facing west. This result is particularly important for many analyses of behavior that are concerned with the behavior itself, rather than where that behavior happens.

One striking feature was DEG’s high performance even on rare behaviors. From our preliminary work building up to the model presented here, we found that simpler models performed well on behaviors that occurred frequently and performed extremely poorly on the infrequent behaviors. Given that in many datasets, the behaviors of interest are infrequent (Fig. 2), we therefore placed a major emphasis on performance in cases with large class imbalances, meaning when some behaviors only occurred in a small fraction of frames. In brief, we accounted for class imbalances in the initialization of the model parameters (Methods). We also changed our cost function to weight errors on rare classes more heavily than errors on common classes. Finally, we tuned the threshold for converting the model’s probability of a given behavior into a classification of whether that behavior was present. Without these added features, the model simply learned to ignore rare classes. We consider these steps toward identifying rare behaviors to be of major significance for effective application in common experimental datasets.

### DeepEthogram accurately predicted behavior bout statistics and transitions

Because DEG produces predictions on individual frames, it allows for subsequent analyses of behavior bouts, such as the number of bouts, the duration of bouts, and the transition probability from one behavior to another. These statistics of bouts are often not available if researchers only record the overall time spent on a behavior with a stopwatch, rather than providing frame-by-frame labels. We found a striking correspondence for the statistics of behavior bouts between the predictions of DEG and those from human labels. On test data, DEG found similar values for the time spent on a given behavior, number of bouts for a behavior in a video, and the mean bout duration (Fig. 4A-F). In addition, the matrix of transition probabilities between behaviors from one frame to the next also had high similarity between DEG predictions and human labels (Fig. 4G). Therefore, the model was able to accurately capture the statistics of the behavioral bouts within videos.

**Figure 4:**
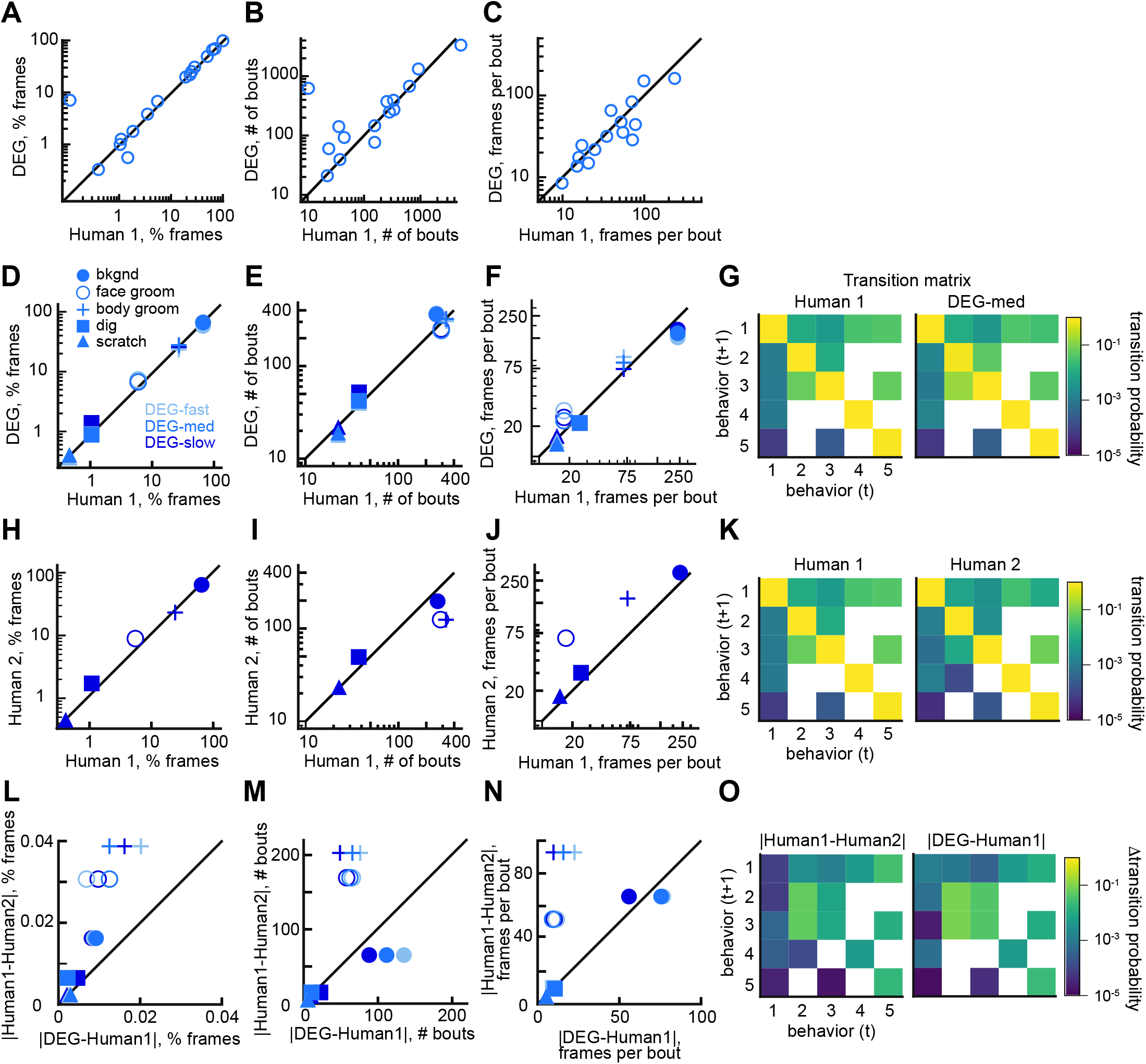
DEG performance on behavior bout statistics. **(A)** Comparison of DEG-medium predictions for the percent of frames with a behavior of interest compared to human labels, using test set data. Each point indicates one behavior across datasets, averaged across three random splits of the data. The off-diagonal point is “defecate” from the mouse-3 dataset. **(B-C)** Similar to (A), except for the number of bouts and the bout length (number of frames per bout). **(D)** Similar to (A), except for behaviors from the Mouse-1 dataset only. All variants of DEG are shown. **(E)** Similar to (B), except for behaviors from the Mouse-1 dataset only. **(F)** Similar to (C), except for behaviors from the Mouse-1 dataset only. **(G)** Transition matrices showing the probability of transition from one behavior on frame i to a behavior on frame i+1 (Methods). **(H)** Comparison of the human labels from Human-1 and Human-2 for the percent of frames with a behavior of interest from the Mouse-1 dataset. **(I-J)** Similar to (H), except for number of bouts and the bout length. **(K)** Similar to (G), except for the two human labelers. **(L)** Comparison of the difference in predictions for the fraction of frames with a given behavior of interest from the Mouse-1 dataset between DEG-medium and Human-1 (x-axis) and Human-1 and Human-2 (y-axis). The higher difference between the human labelers suggests the human labelers had systematic differences in how they labeled behaviors. **(M-N)** Similar to (L), except for the number of bouts and the bout length. **(O)** Comparison of transition matrices between DEG-medium and Human-1 and between Human-1 and Human-2.

To benchmark the model performance on these behavior statistics, we compared the model performance to the level of agreement between two expert human labelers. For the Mouse-1 dataset, which was labeled by multiple researchers, we compared the agreement between human labelers to the agreement between the model predictions and human labels (Fig. 4D-O). In general, the two human labelers had high agreement, showing similar values for the time spent on a behavior, the number of bouts of a behavior, the mean bout duration, and the transition probabilities between behaviors (Fig. 4H-K). Surprisingly, however, the agreement between two human labelers appeared slightly worse than for the model’s performance for particular behaviors, perhaps indicating systematic differences in how two researchers labeled a behavior and thus highlighting potential concerns of inter-human variability and bias (Fig. 4L-O). Given that DEG performed slightly worse on F1 scores relative to expert humans but did just as well as, or maybe better than, expert humans on bout statistics, it is possible that for rare behaviors, DEG may miss a small number of bouts, which would minimally affect bout statistics but could decrease the overall F1 score.

Together, our results from Figures 3-4 and Supplementary Figures 4-6 indicate that DEG’s predictions match well the labels defined by expert human researchers. Further, these model predictions allow easy post-hoc analysis of additional statistics of behaviors, which may be challenging to obtain with traditional manual methods.

### DeepEthogram required little training data to achieve high performance

We evaluated how many video frames a user must label to train a reliable model. We selected 1, 2, 4, 8, 12, or 16 random videos for training and used the remaining videos for evaluation. We only required that each training set had at least one frame of each behavior. We trained the feature extractors, extracted the features, and trained the sequence models for each split of the data. We repeated this process five times for each number of videos, resulting in 30 trained models per dataset. Given the large number of dataset variants for this analysis, to reduce overall computational time, we used DEG-fast and focused on only the Mouse-1, Mouse-2, and Fly datasets. Also, we trained the flow generator only once and kept it fixed for all experiments. For all but the rarest behaviors, the models performed at high levels even with only one labeled video in the training set (Fig. 5A-C). For all the behaviors studied across datasets, the performance measured as accuracy or F1 score approached seemingly asymptotic levels after training on approximately 12 videos. Therefore, a training set of this size or less is likely sufficient for many cases.

**Figure 5:**
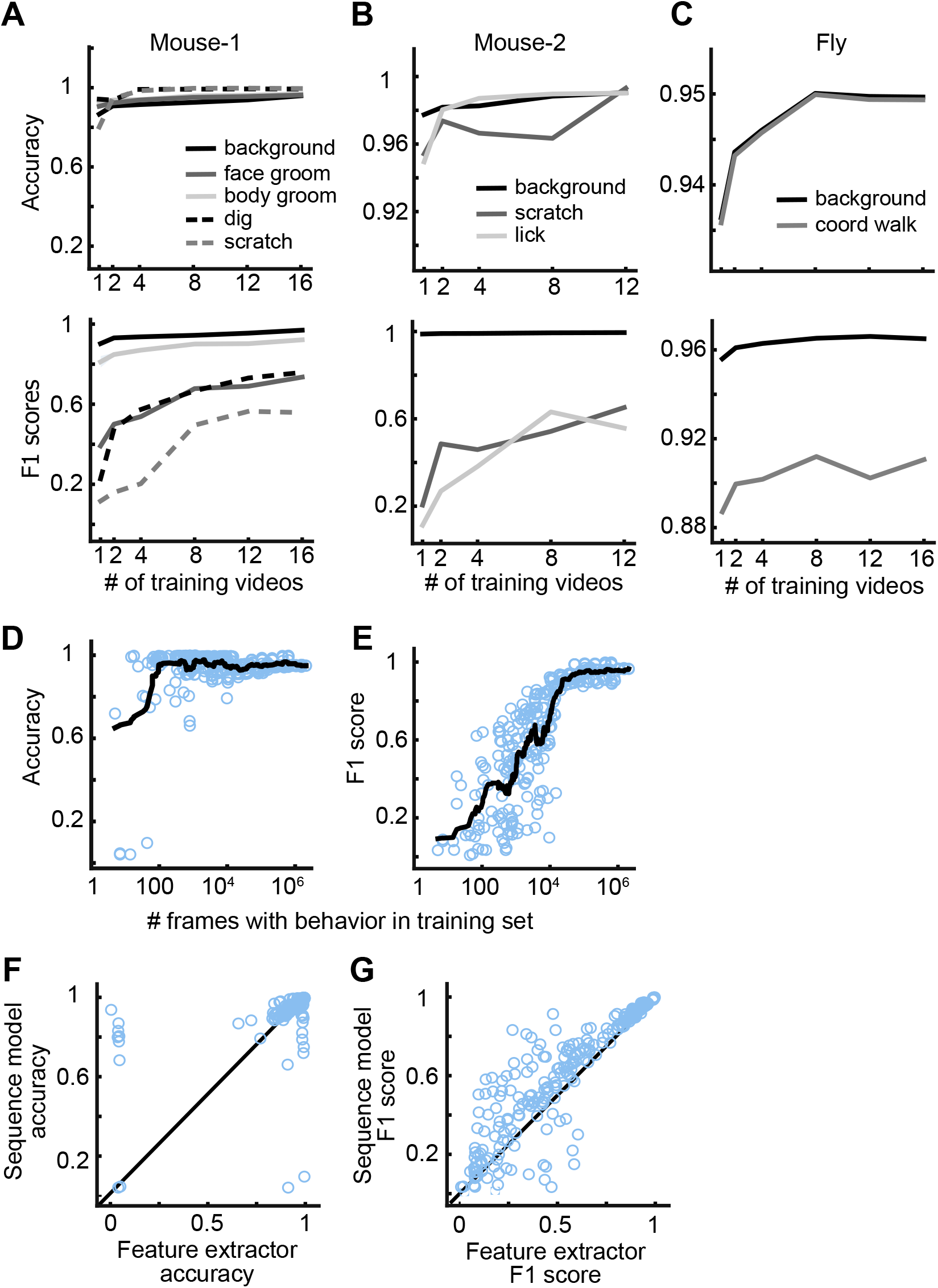
DEG performance as a function of training set size. **(A)** Accuracy (top) and F1 score (bottom) for DEG-fast as a function of the number of videos in the training set for Mouse-1, shown for each behavior separately. The mean is shown across five random selections of training videos. **(B-C)** Similar to (A), except for the Mouse-2 dataset and Fly dataset. **(D)** Accuracy of DEG-fast as a function of the number of frames with the behavior of interest in the training set. Each point is one behavior for one random split of the data, across datasets. The black line shows the running average. For reference, 104 frames is ~5 minutes of behavior at 30 frames per second. **(E)** Similar to (D), except for F1 score. **(F)** Accuracy for the predictions of DEG-fast using the feature extractors only or using the sequence model. Each point is one behavior from one split of the data, across datasets, for the splits used in (D-E). **(G)** Similar to (F), except for F1 score.

We also analyzed the model’s performance as a function of the number of frames of a given behavior present in the training set. For each random split, dataset, and behavior, we had a wide range of the number of frames containing a behavior of interest. Combining all these models together, we found that the model performed with more than 90% accuracy when trained with only 80 example frames of a given behavior and over 95% accuracy with only 100 positive example frames (Fig. 5D). Furthermore, DEG achieved an F1 score of 0.7 with only 9000 positive example frames, which corresponds to about 5 minutes of example behavior at 30 frames per second (Fig. 5E, see Fig. 3J for an example of ~0.7 F1 score). In addition, when the sequence model was used instead of using the predictions directly from the feature extractors, model performance was higher (Fig. 5F-G) and required less training data (data not shown), emphasizing the importance of using long timescale information in the prediction of behaviors. Therefore, DEG models required little training to achieve high performance. As expected, as more training data were added, the performance of the model improved, but this rather light dependency on the amount of training data makes DEG particularly amenable for even smallscale projects. We note that here we used DEG-fast due to the large numbers of splits of the data, and we anticipate that the more complex DEG-medium and DEG-slow models might even require less training data.

### DeepEthogram allows rapid inference time

A key aspect of the functionality of the software is the speed with which the models can be trained and predictions about behaviors made on new videos. Although the versions of DEG varied in speed, they were all fast enough to allow functionality in typical experimental pipelines. On modern computer hardware, the flow generator and feature extractors could be trained in approximately 24 hours. In many cases, these models would only need to be trained once. Afterwards, performing inference to make predictions about the behaviors present on each frame could be performed at ~88 frames per second for videos at 256 x 256 resolution for DEG-fast, at 45 frames per second for DEG-medium, and 15 frames per second for DEG-slow. Thus, for a standard 30-minute video collected at 60 frames per second, inference could be finished in 20 minutes for DEG-fast or two hours for DEG-slow. Importantly, the training of the models and the inference involve zero user time because they do not require manual input or observation from the user. Furthermore, this speed is rapid enough to get results quickly after experiments to allow fast analysis and experimental iteration.

### A graphical user interface for beginning-to-end management of experiments

We developed a graphical user interface (GUI) for labeling videos, training models, and running inference (Fig. 6). To train DEG models, the user first defines which behaviors of interest they would like to detect in their videos. Next, the user imports a few videos into DEG, which will then automatically calculate video statistics and organize them into a consistent file structure. Then the user clicks a button to train the *flow generator* model, which occurs without user time. While this model is training, the user can next go through a set of videos frame-by-frame and label the presence or absence of all behaviors in these videos. Labeling is performed with simple keyboard or mouse clicks at the onset and offset of a given behavior while scrolling through a video in a viewing window. After a small number of videos have been labeled and the *flow generator* is trained, the user then clicks a button to train the *feature extractors*, which occurs without user input and saves the extracted features to disk. Finally, the *sequence model* can be trained automatically on these saved features by clicking another button. All these training steps could in many cases be performed once. With these trained models, the user can import new videos and click the *predict* button, which will estimate the probability of each behavior on each frame. This GUI therefore presents a single interface for labeling videos, training models, and generating predictions on new videos. Importantly, this interface requires no programming by the end-user.

**Figure 6:**
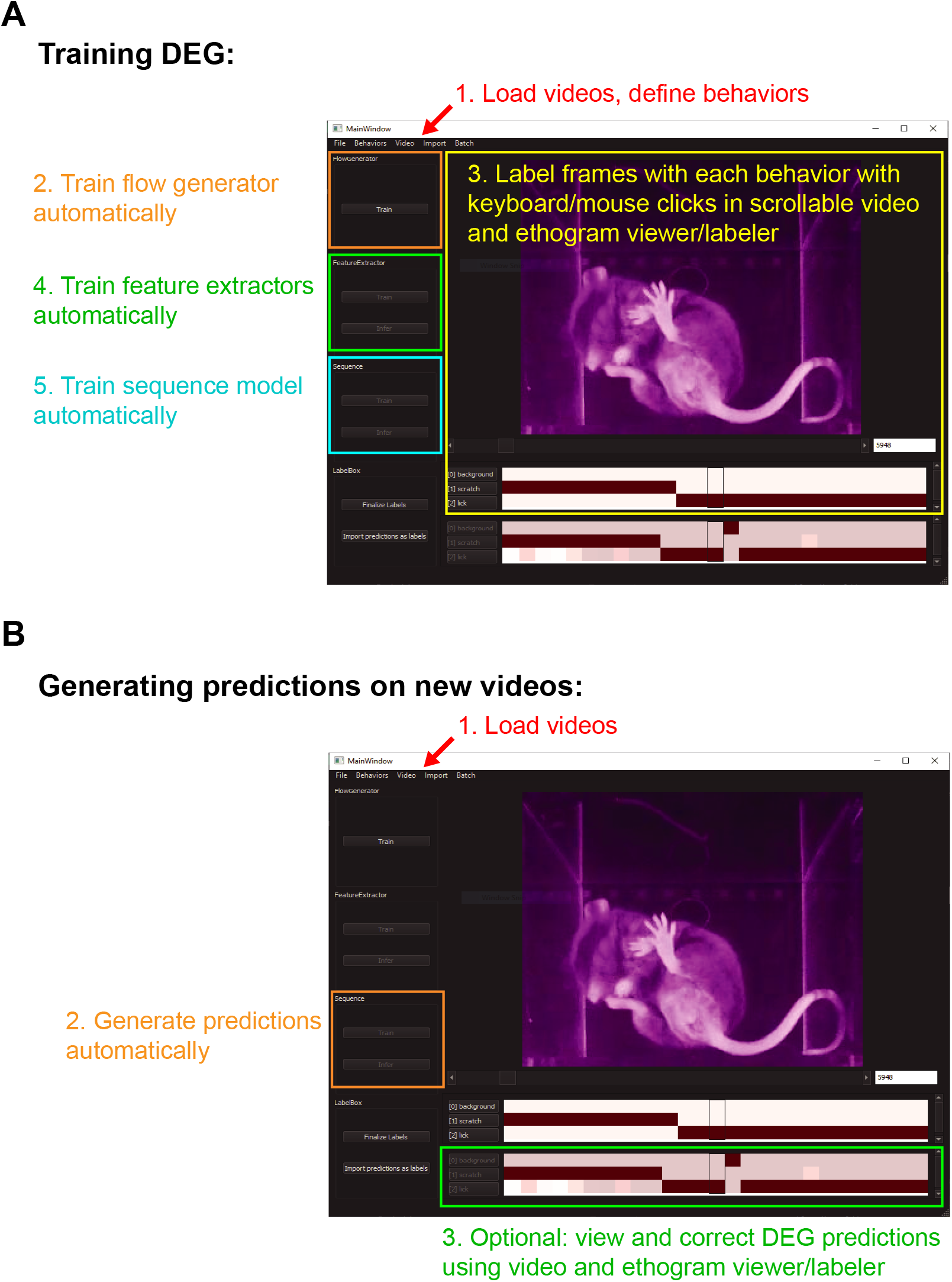
Graphical user interface. **(A)** Example DeepEthogram window with training steps highlighted. **(B)** Example DeepEthogram window with inference steps highlighted.

The GUI also includes an option for users to manually check and edit the predictions output by the model. The user can load into the GUI a video and predictions made by the model. By scrolling through the video, the user can see the predicted behaviors for each frame and update the labels of the behavior manually, just as they would for labeling for the training of the model. This allows users to validate the accuracy of the model and to fix errors should they occur. This process is expected to be fast because the large majority of frames are expected to be labeled correctly, based on our accuracy results, so the user can focus on the small number of frames associated with rare behaviors or behaviors that are challenging to detect automatically. Importantly, these new labels can then be used to re-train the models to obtain better performance on future experimental videos. Documentation for the GUI will be included on the project’s website.

## Discussion

We developed a method for automatically classifying each frame of a video into a set of user-defined behaviors. Our open source software, called DeepEthogram, provides the code and interface necessary to label videos and train DEG models. We show that modern computer vision methods based on pre-trained deep neural networks can be readily applied to animal behavior datasets. DEG performed well on multiple datasets and generalized well across videos and animals, even for identifying extremely rare behaviors. Importantly, by design, CNNs ignore absolute spatial location and thus were able to identify behaviors even when animals were in different locations and orientations within a behavioral arena (Supplementary Fig. 11). We anticipate this software package will save researchers great amounts of time, will lead to more reproducible results by eliminating inter-researcher variability, and will enable experiments that may otherwise not be possible, by increasing the number of experiments a lab can reasonably perform or the number of behaviors that can be investigated. DEG thus joins a growing community of open-source computer vision applications for biological research^8,9,12^.

The models presented here performed well for all datasets tested. In general, we expect the models will perform well in cases in which there is a high degree of agreement between separate human labelers, as our results in Figure 4 indicate. As we have shown, the models do better with more training data. We anticipate that a common use of DEG will be to make automated predictions for each video frame followed by rapid and easy user-based checking and editing of the labels in the GUI for the small number of frames that may be inaccurately labeled. We note that these updated labels can then be used as additional training data to continually update the models and thus improve the performance on subsequent videos. It is important to remember that the models operate directly on the video pixels. This is an advantage because pre-processing is not required, and the researcher does not need to make decisions about which features of the animal to track. Furthermore, the use of raw pixel values allows for a general purpose pipeline that can be applied to videos of all types, without the need to tailor pre-processing steps depending on the behavior of interest, species, number of animals, video angles, resolution, and maze geometries. However, because DEG operates on raw pixels, it is possible that our models may perform more poorly in zoomed-out videos in which the animal is only a few pixels. In addition, it is possible that specialized pipelines designed specifically for one task, such as mouse social behaviors^7^, could perform better at that specific task. An alternate approach to ours is to use innovative methods for estimating pose, including DeepLabCut^22,23^, LEAP^27^, and others^25^, followed by frame-by-frame classification of behaviors based on pose in a supervised^7,19,21^ or unsupervised^17^ way. Using pose for classification could make behavior classifiers faster to train and less susceptible to overfitting. Depending on the design, such skeleton-based action recognition could more easily predict behaviors separately for each of multiple animals in a video, as JAABA does^18^.

DEG may prove to be especially useful when a large number of videos or behaviors need to be analyzed in a given project. These cases could include drug discovery projects or projects in which multiple genotypes need to be compared. Additionally, DEG could be used for standardized behavioral assays, such as those run frequently in a behavioral core facility or across many projects. Importantly, whereas user-time scales linearly with the number of videos for manual labeling of behaviors, user-time for DEG is limited to only the labeling of initial videos for training the models and then can involve essentially no time on the user’s end for all subsequent movies. In our hands, it took approximately 1-3 hours for an expert researcher to label five behaviors in a ten-minute movie from the Mouse-3 dataset. This large amount of time was necessary for researchers to scroll back and forth through a movie to mark behaviors that are challenging to identify by eye. If only approximately 10 human-labeled movies are needed for training the model, then only approximately 10-30 hours of user time would be required. Subsequently, tens to hundreds to thousands of movies could be analyzed, across projects and labs, without additional user-time, which would normally cost additionally hundreds to thousands of hours of time from researchers. Notably, the use of DEG should make results more reproducible across studies and reduce variability imposed by inter-human labeling differences, as we showed in Figure 4. Furthermore, in neuroscience experiments, DEG could aid identification of the starts and stops of movements to relate to neural activity measurements or manipulations. It would be interesting to use DEG’s optic flow snippets as inputs to unsupervised behavior pipelines, where they could help to uncover latent structure in animal behavior^13,15,16^.

Future extensions could continue to improve the accuracy and utility of DEG. First, DEG could be easily combined with an algorithm to track an animal’s location in an environment^2^, thus allowing the identification of behaviors of interest and where those behaviors occur. Also, while the use of CNNs for classification is standard practice in machine learning, recent works in temporal action detection use widely different sequence modeling approaches and loss functions^29,32,39^. Testing these different approaches in the DEG pipeline could further improve performance. Importantly, DEG was designed in a modular way to allow easy incorporation of new approaches as they become available. While inference is already fast, further development could improve inference speed by using low-precision weights, model quantization, or pruning. Furthermore, although our model is currently designed for temporal action localization, DEG could be extended by incorporating models for spatiotemporal action localization, in which there can be multiple actors (i.e., animals) performing different behaviors on each frame.

## Code and Data Availability

Code for DeepEthogram and its GUI is available at: https://github.com/jbohnslav/deepethogram. We will make datasets available upon reasonable request to the corresponding author.

## Acknowledgements

We thank Woolf lab members Rachel Moon for data acquisition and scoring and Victor Fattori for scoring, and David Roberson and Lee Barrett for designing and constructing the PalmReader and iBob mouse viewing platforms. We thank the Harvey lab for helpful discussions and feedback on the manuscript. This work was supported by grants from the NIH (R01MH107620 (CDH), R01NS089521 (CDH), R01NS108410 (CDH), F31NS108450 (JPB), R35NS105076 (CJW), R00NS101057 (LLO), K99DE028360), the European Research Council (ERC-Stg-759782 (EC)), an NSF GRFP (NKW), a Harvard Medical School Dean’s Innovation Award (CDH), and a Harvard Medical School Goldenson Research Award (CDH).

## Author Contributions

JPB, NKW, CJW, CDH conceived of the project. JPB and CDH developed the approach. JPB developed the software and analyzed the data. NKW, KJC, DY, and TC performed experiments. NKW, KJC, DY, TC, and JPB labeled the videos. EC, LLO, and CJW supervised the experiments. CDH supervised the software development and data analysis. JPB and CDH wrote the manuscript with input from all authors.

## Methods

### DeepEthogram pipeline

Along with this publication, we are releasing open source Python code for labeling videos, training all DeepEthogram models, and performing inference on new videos. The code, associated documentation, and files for the GUI can be found at https://github.com/jbohnslav/deepethogram.

#### Implementation

We implemented DeepEthogram in the Python programming language (version 3.7 or later)^40^. We used PyTorch^41^ (version 1.4.0) for all deep learning models. We used OpenCV^42^ for image reading, writing, and image augmentations. We used scikit-learn^43^ for evaluation metrics, along with custom Python code. We used Nvidia GeForce 1080 Ti GPUs for all models except pretrained Kinetics700 models, for which we used either the 1080 Ti or a Titan RTX. Inference speed was evaluated on a computer running Ubuntu 18.04, an AMD Ryzen Threadripper 2950X CPU, an Nvidia Geforce 1080 Ti, a Samsung 970 Evo hard disk, and 128 GB DDR4 memory.

### Datasets

All experimental procedures were approved by the Institutional Animal Care and Use Committees at Boston Children’s Hospital or Massachusetts General Hospital and were performed in compliance with the Guide for the Care and Use of Laboratory Animals.

### Kinetics700

To pretrain our models for transfer to neuroscience datasets, we use the Kinetics700^31^ dataset. The training split of this dataset consisted of 538,523 videos and 141,677,361 frames. We first resized each video so that the short side was 256 pixels. During training, we randomly cropped 224 x 224 pixel images, and during validation, we used the center crop.

### Mouse-1

Recordings of voluntary behavior were acquired for 14 adult male C57/B6J mice on the PalmReader device (Roberson et al., submitted). In brief, images were collected with infrared illumination and frustrated total internal reflectance (FTIR) illumination on alternate frames. The FTIR channel highlighted the parts of the mouse’s body that were in contact with the floor. We stacked these channels into an RGB frame: red corresponded to the FTIR image, green corresponded to the infrared image, and blue was the pixel-wise mean of the two. In particular, images were captured as a ventral view of mice placed within an opaque 18 cm long x 18 cm wide x 15 cm high chamber with a 5 mm thick borosilicate glass floor using a Basler acA2000-50gmNIR GigE near-infrared camera at 25 frames per second. Animals were illuminated from below using non-visible 850 nm near-infrared LED strips. All mice were habituated to investigator handling in short (~5 minute) sessions and then habituated to the recording chamber in 2 sessions lasting 2 hours on separate days. On recording days, mice were habituated in a mock recording chamber for 45 minutes, and then moved by an investigator to the recording chamber for 30 minutes. Each mouse was recorded in 2 of these sessions spaced 72 hours apart. The last 10 minutes of each recording was manually scored on a frame-by-frame basis for defined actions using a custom interface implemented in MATLAB. The 28 ~10-minute videos totaled 419,846 frames (and labels) in the dataset. Data were recorded at 1000 x 1000 pixels and down-sampled to 250 x 250 pixels. We resized to 256 x 256 pixels using bilinear interpolation during training and inference.

For human-human comparison, we re-labeled all videos using the DeepEthogram GUI. Previous labels were not accessible during re-labeling. Criterion for re-labeling were written in detail by the original experimenters, and example labeled videos were viewed extensively before re-labeling.

### Mouse-2

Recordings of voluntary behavior were acquired for 16 adult male and female C57/B6J mice. These data were collected on the iBob device (Roberson et al., submitted). Briefly, the animals were enclosed in a device containing an opaque six-chambered plastic enclosure atop a glass floor. The box was dark and illuminated with only infrared light. Animals were habituated for one hour in the device before being removed to clean the enclosure. They were then habituated for another 30 minutes and recorded for 30 minutes. Recorded mice were either wild type or contained a genetic mutation predisposing them to dermatitis; thus scratching and licking behavior were scored. Up to six mice were imaged from below simultaneously and subsequently cropped to a resolution of 270 x 240 pixels. Images were resized to 256 x 256 pixels during training and inference. Data were collected at 30 frames per second. There were 16 ~30-minute videos for a total of 863,232 frames.

### Mouse-3

Videos for the mouse-3 dataset were obtained from published studies^44,45^ and unpublished work (Clausing et al., unpublished). Video recordings of voluntary behavior were acquired for 20 adult male mice.

All mice were exposed to a novel empty arena (40 cm x 40 cm x 40 cm) with opaque plexiglass walls. Animals were allowed to explore the arena for 10 minutes, under dim lighting. Videos were recorded via an overhead-mounted camera at either 30 or 60 frames per second. Videos were acquired with 2-4 mice simultaneously in separate arenas and cropped with a custom python script such that each video contained the behavioral arena for a single animal. Prior to analysis, some videos were brightened in FIJI^46^, using empirically determined display range cutoffs that maximized the contrast between the mouse’s body and the walls of the arena. Twenty of the 10-minute recordings were manually scored on a frame-by-frame basis for defined actions in the DeepEthogram interface. All data were labeled by an experimenter. The twenty ~10 minute videos totaled 537,534 frames (and labels).

For human-human comparison, we re-labeled a subset of 5 videos using the DeepEthogram GUI. Previous labels were not accessible during re-labeling. Criterion for re-labeling were written in detail by the original experimenters, and example labeled videos were viewed extensively before re-labeling.

### Fly

Wild Type DL adult male flies (Drosophila melanogaster), 2–4 days post-eclosion were reared on a standard fly medium and kept on a 12-h light-dark cycle at 25°. Flies were cold anesthetized and placed in a fly sarcophagus. We glued the fly head to its thorax and finally to a tungsten wire at an angle around 60 degrees (UV cured glue, Bondic). The wire was placed in a micromanipulator used to position the fly on top of an air-suspended ball. Side-view images of the fly were collected at 200 Hz with a Basler A602f camera. Videos were down-sampled to 100 Hz. There were 19 ~30 minute videos for a total of 3,419,943 labeled frames. Images were acquired at 168 x 100 pixels and up-sampled to 192 x 128 pixels during training and inference. Images were acquired in grayscale but converted to RGB (cv2.cvtColor, cv2.COLOR_GRAY2RGB) so that input channels were compatible with pretrained networks and other datasets.

## Models

### Overall setup

Problem statement: Our input features were a set of images with dimensions [T, C, H, W], and our goal was to output the probability of each behavior on each frame, which is a matrix with dimensions [T, K]. *T* is the number of frames in a video. *C* is the number of input channels—in typical color images, this number is three, for the red, green, and blue (RGB) channels. *H, W* are the height and width of the images in pixels. *K* is the number of user-defined behaviors we aimed to estimate from our data.

#### Training protocol

We used the ADAM optimizer^47^ with an initial learning rate of 1 x 10^-4^ for all models. When validation performance saturated for 10,000 training steps, we decreased the learning rate by a factor of 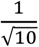 on the Kinetics700 dataset, or by a factor of 0.1 for neuroscience datasets (for speed). We stopped training when the learning rate dropped below 5*e* – 7. For Kinetics700^31^, this required about 800,000 training steps. For feature extractors in neuroscience datasets, this typically required 100,000 training steps. For Kinetics700, we used the provided train and validation split. For neuroscience datasets, we randomly picked 60% of videos for training, 20% for validation, and 20% for test (with the exception of the subsampling experiments for Figure 5, wherein we only used training and validation sets to reduce overall training time). Our only restriction on random splitting was ensuring that at least one frame of each class was included in each split. We were limited to 3 random splits of the data for most experiments due to the computational and time demands of retraining models.

#### End-to-end training

We could, in theory, train the entire DEG pipeline end-to-end. However, we chose to train the flow generator, feature extractor, and then sequence models sequentially. By backpropagating the classification loss into the flow generator^28^, we risk increasing the overall number of parameters and the risk of overfitting. Furthermore, we designed the sequence models to have a large temporal receptive window. We therefore train on long sequences (see below). Very long sequences of raw video frames take large amounts of VRAM and exceed our computational limits. By illustration, to train on sequences of, for example, 180 snippets of 11 images, our tensor would be of shape [N x 33 x 180 x 256 x 256]. This corresponds to 24 GB of VRAM at a batch size of 16, just for the data and none of the neural activations or gradient, which is impractical. Therefore, we first extract features to disk and subsequently train sequence models.

#### Augmentations

To improve the robustness and generalization of our models, we augmented the input images with random perturbations for all datasets. We perturbed the image brightness and contrast, rotated each image by up to 10 degrees, and randomly flipped each image horizontally with a probability of 0.5. The input to the flow generator model is a set of 11 frames; the same augmentations were performed on each image in this stack. On Mouse-1-3, we also flipped images vertically with a probability of 0.5. We calculated the mean and standard deviation of the RGB input channels and standardized the input channel-wise.

#### Pretraining + transfer learning

All flow generators and feature extractors were first trained to classify videos in the Kinetics700 dataset (see below). These weights were used to initialize models on neuroscience datasets. Sequence models were trained from scratch.

### Flow generators

For optical flow extraction, a common algorithm to use is TV-L1^48^. However, common implementations of this algorithm^42^ require compilation of C++, which would introduce many dependencies and make installation more difficult. Furthermore, recent work^28^ has shown that even simple neural-network-based optical flow estimators outperform TV-L1 for action detection. Therefore, we used CNN-based optical flow estimators. Furthermore, we found that saving optic flow as JPEG images, as is common, significantly degraded performance. Therefore, we computed optic flows from a stack of RGB images at runtime for both training and inference. This method is known as Hidden Two-Stream Networks^28^.

#### Architectures

##### TinyMotionNet

For every timepoint, we extracted features based on one RGB image and up to 10 optical flow frames (see Two Stream section). Furthermore, for large datasets like Kinetics700^31^, it was time-consuming and required a large amount of disk space to extract and save optical flow frames. Therefore, we implemented TinyMotionNet^28^ to extract 10 optical flow frames from 11 RGB images “on the fly”, as we extracted features. TinyMotionNet is a small and fast optical flow model with 1.9 million parameters. Similar to a U-Net^49^, it consists of a downward branch of convolutional layers of decreasing resolution and increasing depth. It is followed by an upward branch of increasing resolution. Units from the downward branch are concatenated to the upward branch. During training, estimated optic flows were output at 0.5, 0.25, and 0.125 of the original resolution.

##### MotionNet

MotionNet is similar to TinyMotionNet except with more parameters and more feature maps per layer. During training, estimated optic flows were output at 0.5, 0.25, and 0.125 of the original resolution. See the original paper^28^ for more details.

##### TinyMotionNet3D

This novel architecture is based on TinyMotionNet^28^, except we replaced all 2D convolutions with 3D convolutions. We maintained the height and width of the kernels. On the encoder and decoder branches, we used a temporal kernel size of three, meaning that each filter spanned three images. On the last layer of the encoder and the *iconv* layers that connect the encoder and decoder branches, we used a temporal kernel of two, meaning the kernels spanned two consecutive images. We aimed to have the model learn the displacement between two consecutive images (that is, the optic flow). Due to the large memory requirements of 3D convolutional layers, we used 16, 32, and 64 filter maps per layer. For this architecture, we noticed large estimated flows in texture-less regions in neuroscience datasets after training on Kinetics. Therefore, we added a L1 sparsity penalty on the flows themselves (see loss functions, below).

##### Modifications

For the above models, we deviated from the original paper. First, each time the flows were up-sampled by a factor of two, we multiplied the values of the neural activations by two. If the flow size increased from 0.25 to 0.5 of the original resolution, a flow estimate of 1 corresponds to four pixels and two pixels in the original image, respectively. To compensate for this distortion, we multiplied the up-sampled activations by two. Secondly, when used in combination with the CNN feature extractors (see below), we did not compress the flow values to the discrete values between 0 and 255^28^. In fact, we saw performance increases for keeping the continuous float32 values. Third, we did not backpropagate the classifier loss function into the flow generators, as the neuroscience datasets likely did not have enough training examples to make this a sensible strategy. Finally, for MotionNet, we only output flows at three resolutions (rather than five) for consistency.

#### Loss functions

In brief, we train flow generators to minimize reconstruction errors and minimize high-frequency components (to encourage smooth flow outputs).

#### MotionNet Loss

For full details, see original paper^28^. For clarity, we reproduce the loss functions here. We estimate the current frame given the next frame and an estimated optic flow as follows:

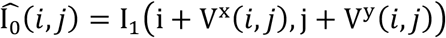

*I*_0_, *I*_1_ are the current and next image. *i, j* are the indices of the given pixel in rows and columns.V^x^(*i, j*), V^y^(*i, j*) are the estimated x and y displacements between *I*_0_, *I*_1_, which means *V* is the optic flow. We use Spatial Transformer Networks^50^ to perform this sampling operation in a differentiable manner (PyTorch function *torch.nn.functional.grid_sample*).

The image loss is the error between the reconstructed 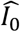 and original *I*_0_.

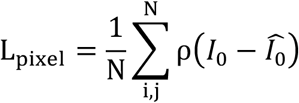

*ρ* is the generalized Charbonnier penalty *ρ*(*x*) = (*x*^2^ + *ϵ*^2^)^*α*^, which reduces the influence of outliers compared to a simple L1 loss. Following Zhu et al.^28^, we use *α* = 0.4, *ϵ* = 1*e*^−7^.

The structural similarity (SSIM^51^) loss encourages the reconstructed 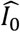 and original *I*_0_ to be perceptually similar:

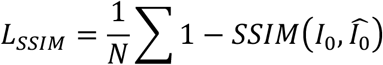

The smoothness loss encourages smooth flow estimates by penalizing the *x* and *y* gradients of the optic flow:

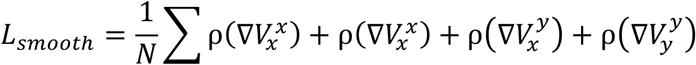

We set the Charbonnier α = 0.3.

For the TinyMotionNet3D architecture only, we added a flow sparsity loss that penalizes unnecessary flows:

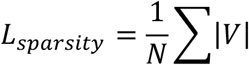

The total loss is the weighted sum of the previous components:

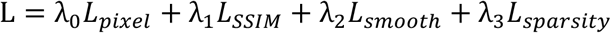

Following Zhu et al.^28^, we set λ_0_ = 1, λ_1_ = 1. During training, the flow generators output flows at multiple resolutions. From largest to smallest, we set λ_2_ to be 0.01, 0.02, 0.04. For TinyMotionNet3D, we set λ_3_ to 0.05 and reduced λ_2_ by a factor of 0.25.

### Feature extractors

The goal of the feature extractor was to model the probability that each behavior was present in the given frame of the video (or optic flow stack). We used two-stream convolutional neural networks^28,52^ to classify inputs from both RGB frames and optic flow frames. These CNNs reduced an input tensor from (*N,C,H,W*) pixels to (*N*, 512) features. Our final fully-connected layer estimated probabilities for each behavior, with output shape (*N,K*). Here, *N* is the batch size. We trained these CNNs on our labels, and then used the penultimate (*N*, 512) *spatial features* or *flow features* as inputs to our sequence models (below).

#### Architectures

We used the ResNet^36,37^ family of models for our feature extractors, one for the spatial stream and one for the flow stream. For DEG-fast, we used ResNet18 with ~11 million parameters. For DEG-medium we used a ResNet50 with ~23 million parameters. We added dropout^53^ layers before the final fully connected layer. For DEG-medium, we added an extra fully-connected layer of shape (2048,512) after the global average pooling layer to reduce the file size of stored features. For DEG-slow, we used a 3D ResNet34^37^ with ~63 million parameters. For DEG-fast and DEG-medium, these models were pre-trained on ImageNet^30^ with three input channels (RGB). We stacked ten optic flow frames, for 20 input channels. To leverage ImageNet weights with this new number of channels, we used the mean weight across all three RGB channels and replicated it 20 times^54^.

#### Loss functions

Our problem is a multi-label classification task. Each timepoint can have multiple positive examples. For example, if a mouse is licking its forepaw and scratching itself with its hindlimb, both “lick” and “scratch” should be positive. Therefore, we used a weighted binary cross entropy loss between our labels and our outputs (PyTorch function *torch.nn.BCEWithLogitsLoss*):

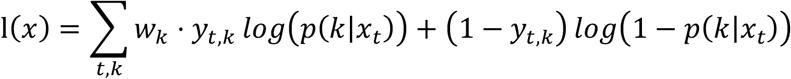

*y_t,k_* is the ground truth label and was 1 if class *k* occurred at time *t*, or otherwise was 0. *p*(*k|x_t_*) is our model output for class *k* at time *t*. Note that for the feature extractor, we only considered one timepoint at a time, so *t* = 0. *w_k_* is a weight given to positive examples—note that there was no corresponding weight in the second term when our ground truth is 0. Intuitively, if we had a very rare behavior, we wanted to penalize the model more for an error on positive examples because there were so few examples of the behavior. We calculated the weight as follows:

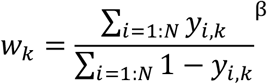

The numerator is the total number of positive examples in our training set, and the denominator is the total number of negative examples in our training set. β is a hyperparameter that we tuned manually. If β = 1, positive examples were weighted fully by their frequency in the training set. If β = 0, all training examples were weighted equally. By illustration, if only 1% of our training set had a positive example for a given behavior, with β = 1 our weight was 100 and with β = 0 this *w_k_* = 1. We empirically found that with rare behaviors β = 1 drastically increased the levels of false positives, while with β = 0 many false negatives occurred. For the Fly dataset with little class imbalance (see Figure 2), we set β = 1 while for Mouse-1, Mouse-2, and Mouse-3 datasets, we set β = 0.5. This *w_k_* argument corresponds to *pos_weight* in *torch.nn.BCEWithLogitsLoss*.

#### Bias initialization

To combat the effects of class imbalance, we set the bias parameters on the final layer to approximate the class imbalance (https://www.tensorflow.org/tutorials/structured_data/imbalanced_data). For example, if we had 99 negative examples and 1 positive example, we wanted to set our initial biases such that the model guessed “positive” around 1% of the time. Therefore, we initialized the bias term as the log ratio of positive examples to negative examples:

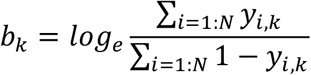

#### Fusion

There are many ways to fuse together the outputs of the spatial and motion stream in two-stream CNNs^55^. For simplicity, we used late, average fusion. We averaged together the 512 dimensional output vectors of the CNNs before the sigmoid function:

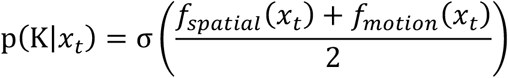

#### Curriculum training

To train the feature extractors, we could have trained the entire model simultaneously end-to-end. We first trained the spatial stream to convergence, followed by the flow stream, and finally we trained the spatial and flow streams simultaneously. We reset the learning rate after each model finished training.

### Sequence models

#### Architecture

The goal of the sequence model was to have a wide temporal receptive field for classifying timepoints into behaviors. For human labelers, it is much easier to classify the behavior at time *t* by watching a short clip centered at *t* rather than viewing the static image. Therefore, we used a sequence model that takes as input a sequence of *spatial features* and *flow features* output by the feature extractors. Our criteria were to find a model that had a large temporal receptive field, as context can be useful for classifying frames. However, we also wanted a model that had relatively few parameters, as this model was trained from scratch on small neuroscience datasets. Therefore, we chose Temporal Gaussian Mixture^29^ (TGM) models, which are designed for temporal action detection. Unless otherwise noted, we used the following hyperparameters:

- Filter length *L* = 15
- Number of input layers *C_in_* = 1
- Number of output layers *C_out_* = 1
- Input dropout = 0.4
- Dropout of output features = 0.5
- Input dimensionality (concatenation of flow and spatial) *D* = 1024
- Number of filters = 8
- Sequence length = 180
- 1D-convolution, not soft attention
- We do not use super-events

For more details, see the Piergiovanni and Ryoo.^29^

#### Modifications

TGM models use two main features to make the final prediction: the [*T, D*] input features (in our case, spatial and flow features from the feature extractors); and the [*T, D*] learned features output by the TGM layers. The original TGM model performed “early fusion” by concatenating these two features into shape [*T*, 2*D*] before the 1D convolution layer. We found in low-data regimes that the model ignored the learned features, and therefore reduced to a simple 1D convolution. Therefore, we performed “late fusion”—we used separate 1D convolutions on the input features and on the learned features. We averaged the output of these two layers (both [*T, K*] activations before the sigmoid function). Secondly, in the original TGM paper, the penultimate layer was a standard 1D convolutional layer with 512 output channels. We found this dramatically increased the number of parameters without improving performance significantly. Therefore, we removed this layer. The final number of parameters is ~12,000.

#### Loss function

We used a weighted binary cross entropy loss (see *feature extractor loss function* above).

**Table 1:** Model summary.

| Model name | Flow generator (parameters) | Feature extractor (parameters) | Sequence model (parameters) | # frames input to flow generator | # frames input to RGB feature extractor | Total params |
| --- | --- | --- | --- | --- | --- | --- |
| DEG-fast | TinyMotionNet (1.9M) | ResNet18 x 2 (22.4M) | TGM (12K) | 11 | 1 | ~ 24.3M |
| DEG-medium | MotionNet (45.8M) | ResNet50 x 2 (49.2M) | TGM (12K) | 11 | 1 | ~ 95M |
| DEG-slow | TinyMotionNet3D (0.4M) | ResNet3D-34 x 2 (127M) | TGM (12K) | 11 | 11 | ~ 127.4M |

### Postprocessing

The output of the feature extractor and sequence model is the probability of behavior *k* occurring on frame *t*: *p_t,k_* = *f*(*x_t,k_*). To convert these probabilities into binary predictions, we thresholded the probabilities:

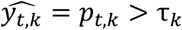

We picked the threshold τ_*k*_ for each behavior *k* that maximized the F1 score (below). We picked the threshold independently on the training and validation sets. On test data, we used the validation thresholds.

We found that these predictions overestimated the overall number of bouts. In particular, very short bouts were over-represented in model predictions. For each behavior *k*, we removed both “positive” and “negative” bouts (binary sequences of 1s and 0s, respectively) of less than 2 frames long.

Finally, we computed the “background” class as the logical not of the other predictions.

### Evaluation and metrics

We used the following metrics: overall accuracy, F1 score, and the area under the receiver operating curve (AUROC) by class. Accuracy was defined as:

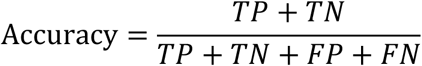

Where *TP* is the number of true positives, *TN* is the number of true negatives, *FP* is the number of false positives, and *FN* is the number of false negatives. We reported overall accuracy, not accuracy for each class.

F1 score was defined as:

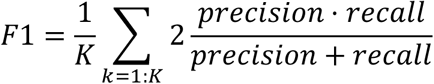

Where 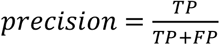 and 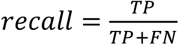. The above was implemented by *sklearn.metrics.f1_score* with argument *average=‘macro’*.

AUROC was computed by taking the area under the receiver operating curve for each class and averaging the result. This was implemented by *sklearn.metrics.roc_auc_score* with argument *average=‘macro’*.

## SUPPLEMENTARY FIGURES

**Supplementary Figure 1:**
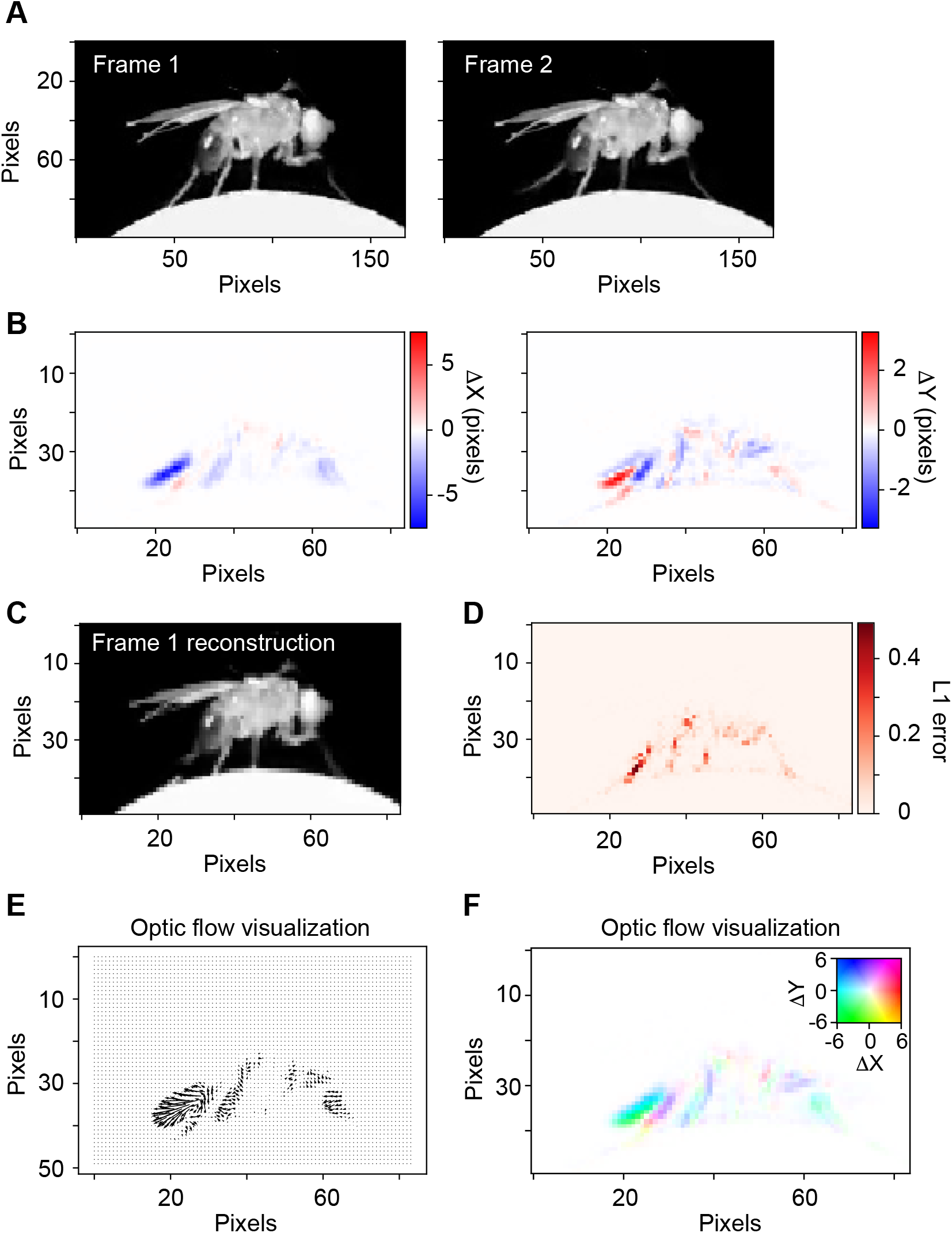
Optic flow. **(A)** Example images from the Fly dataset on two consecutive frames. **(B)** Optic flow estimated with TinyMotionNet. Note the image size is half the original due to the TinyMotionNet architecture. Displacements in the x dimension (left) and y dimension (right) between the frames in (A). **(C)** The reconstruction of frame 1 estimated by sampling frame 2 according to the optic flow calculation. The image was resized with bilinear interpolation before resampling. **(D)** Absolute error between frame 1 and the frame 1 reconstructed from optic flow. **(E)** Visualization of optic flow using arrow lengths to indicate the direction and magnitude flow. **(F)** Visualization of optic flow using coloring according the inset color scale. Left displacements are mapped to cyan, right displacements to red, and so on. Saturation indicates the magnitude of displacement.

**Supplementary Figure 2:**
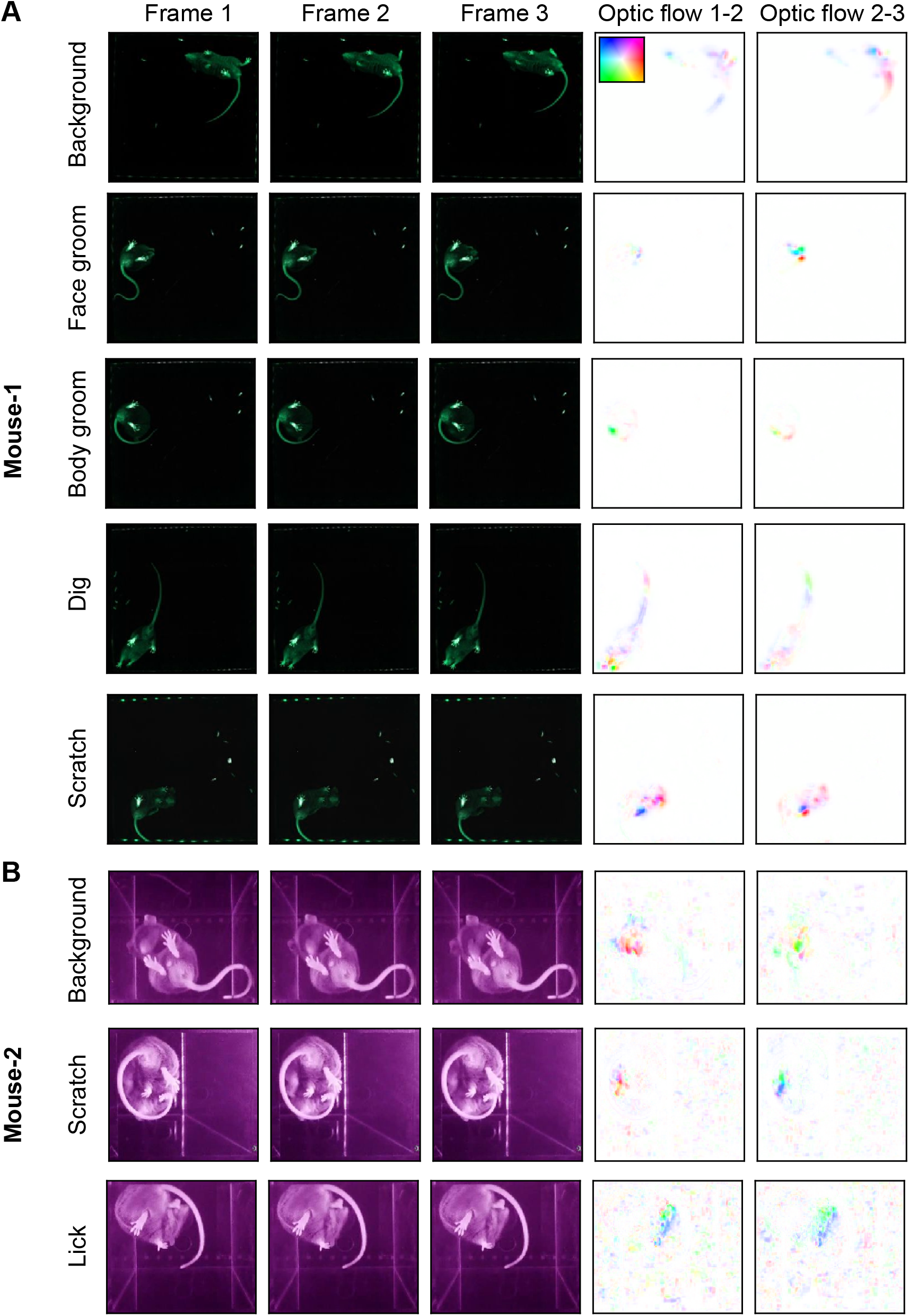
Example images from the datasets, part 1. **(A)** Examples from the Mouse-1 dataset. Each row is three consecutive frames of the indicated behavior. Right columns, optic flow computed by TinyMotionNet and visualized as in Supplementary Fig. 1F. **(B)** Similar to (A), except for the Mouse-2 dataset.

**Supplementary Figure 3:**
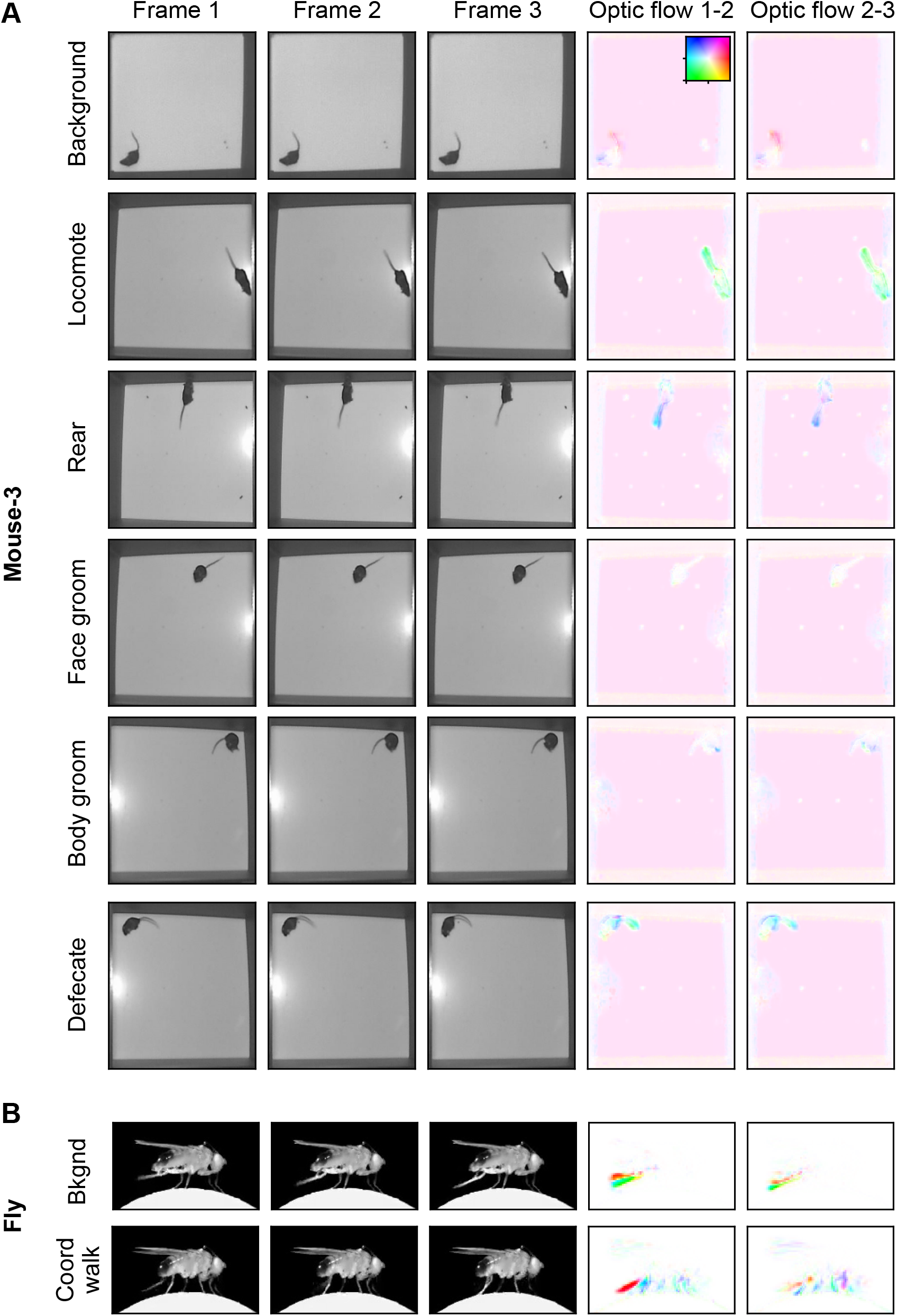
Example images from the datasets, part 2. **(A)** Examples from the Mouse-3 dataset. Each row is three consecutive frames of the indicated behavior. Right columns, optic flow computed by TinyMotionNet and visualized as in Supplementary Fig. 1F. **(B)** Similar to (A), except for the Fly dataset.

**Supplementary Figure 4:**
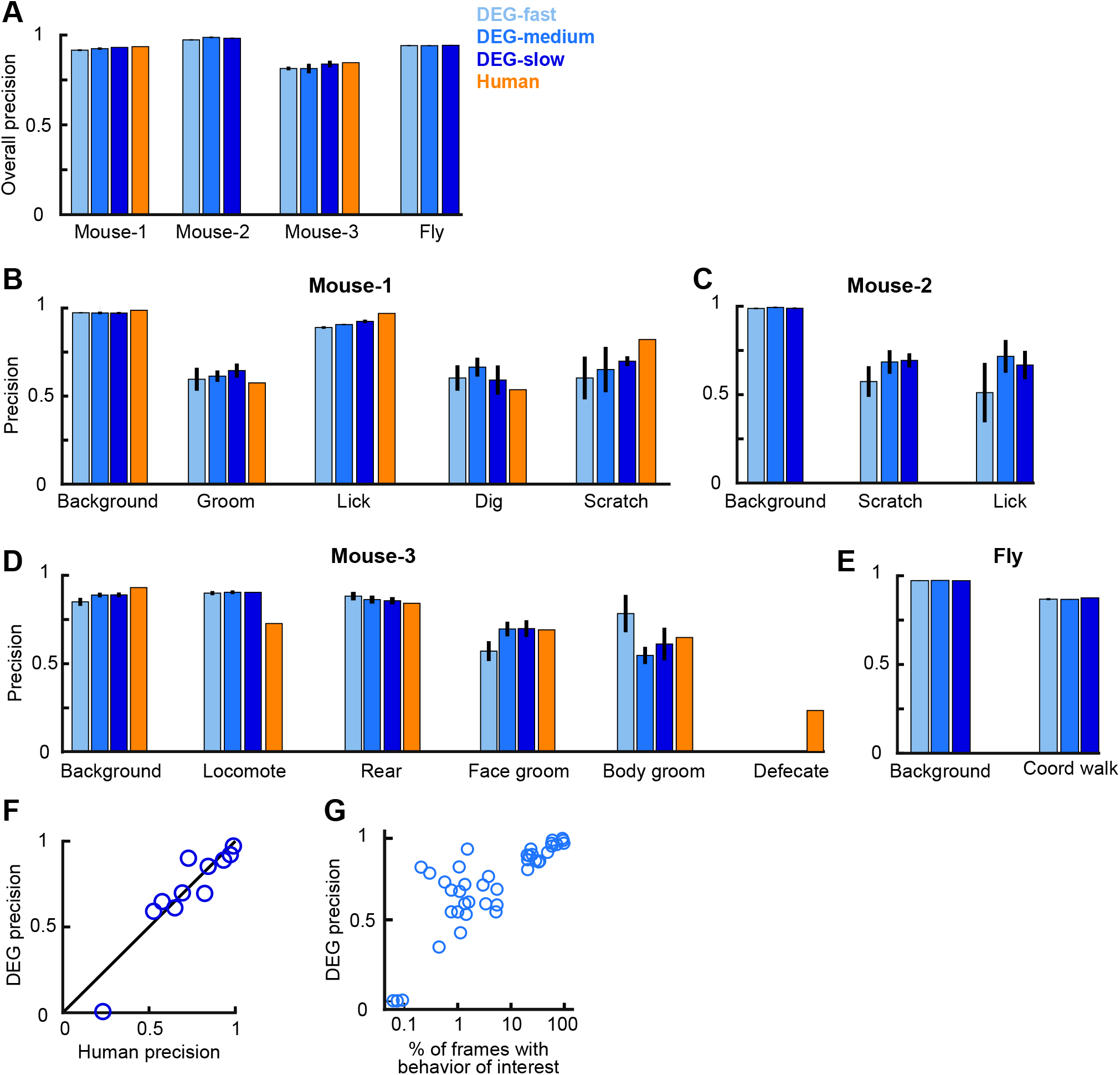
DEG performance measured as precision. **(A)** Overall precision for each DEG variant and dataset for data from the test set. Precision is defined as True Positives / (True Positives + False Positives). Human accuracy corresponds to the agreement between human labelers. Error bars indicate mean +/- s.e.m. across three random splits of the data. **(B)** Precision for each DEG variant for individual behaviors from the Mouse-1 dataset. **(C-E)** Similar to (B), except for the Mouse-2 dataset, Mouse-3 dataset, Fly dataset. **(F)** Precision between two human labelers compared to DEG-slow. Each point is one behavior, with performance averaged across 3 random splits of the data. The point with low DEG precision is for “defecation” from Mouse-3. **(G)** DEG-medium performance measured as precision as a function of the percent of frames containing the behavior of interest in the training data. Each point is for one behavior for one split of the data, with three splits in total.

**Supplementary Figure 5:**
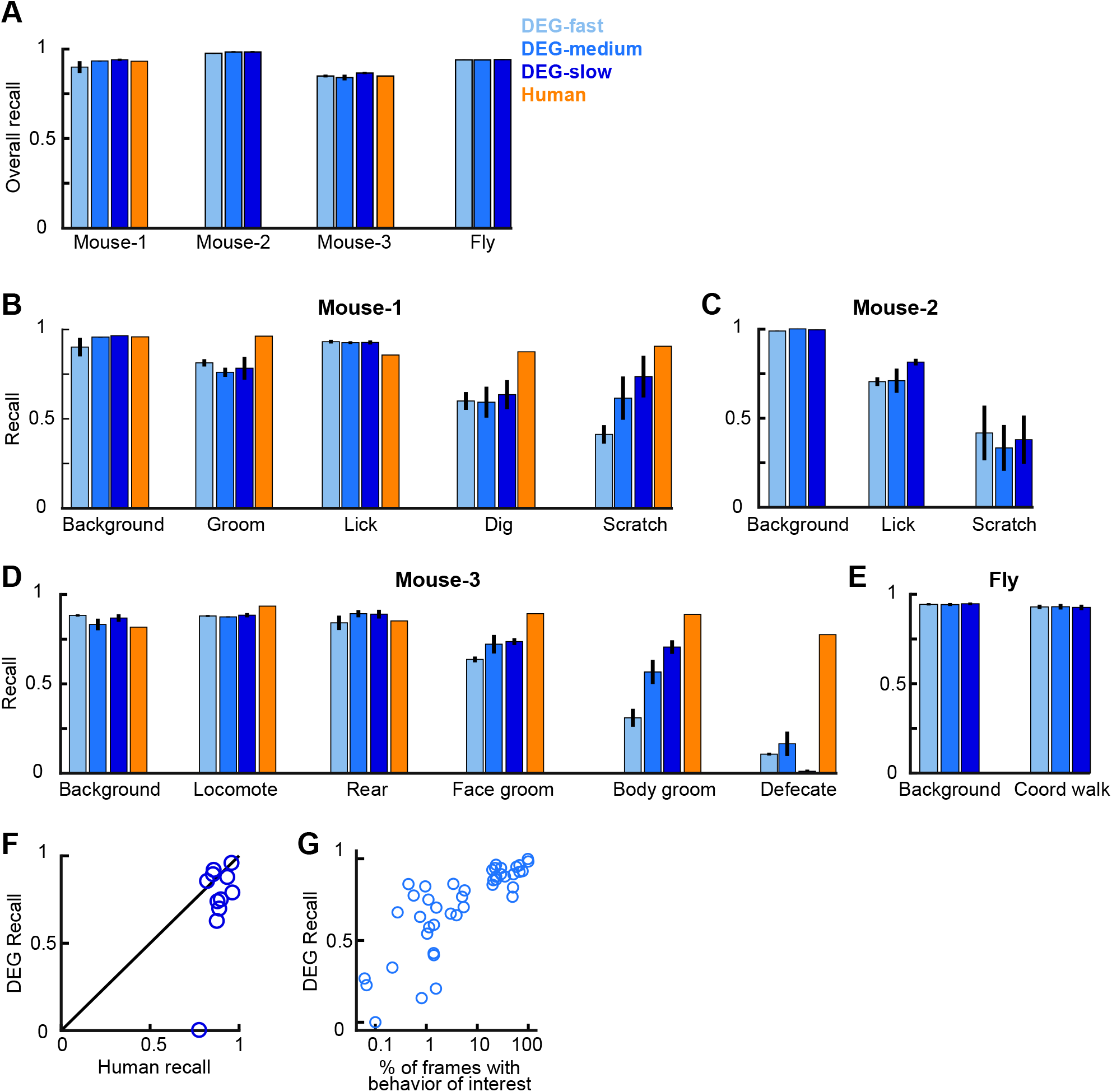
DEG performance measured as recall. **(A)** Overall recall for each DEG variant and dataset for data from the test set. Recall is defined as True Positives / (True Positives + False Negatives). Human accuracy corresponds to the agreement between human labelers. Error bars indicate mean +/- s.e.m. across three random splits of the data. **(B)** Recall for each DEG variant for individual behaviors from the Mouse-1 dataset. **(C-E)** Similar to (B), except for the Mouse-2 dataset, Mouse-3 dataset, Fly dataset. **(F)** Recall between two human labelers compared to DEG-slow. Each point is one behavior, with performance averaged across 3 random splits of the data. The point with low DEG recall is for “defecate” from Mouse-3. **(G)** DEG-medium performance measured as recall as a function of the percent of frames containing the behavior of interest in the training data. Each point is for one behavior for one split of the data, with three splits in total.

**Supplementary Figure 6:**
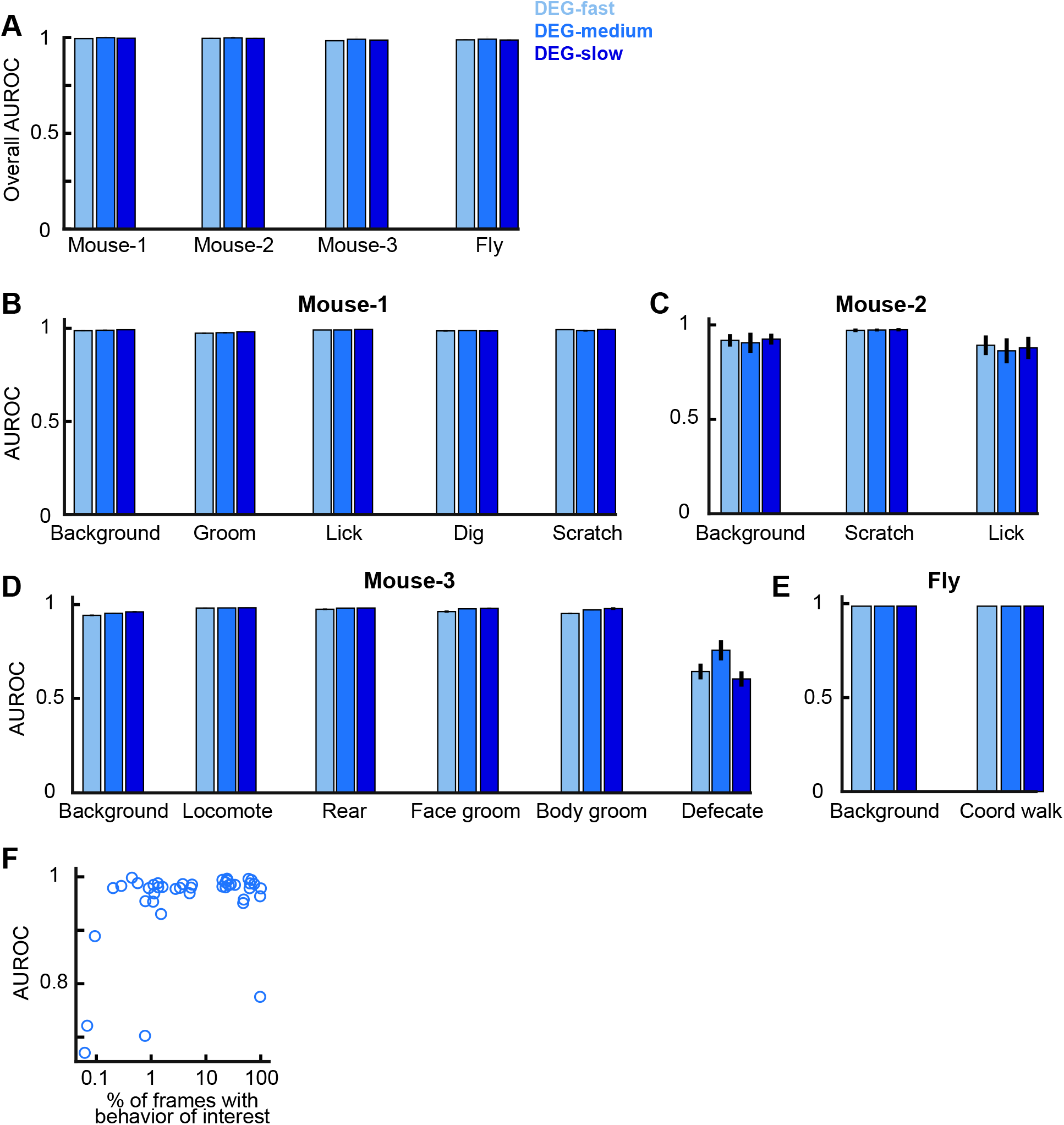
DEG performance measured as AUROC. **(A)** Overall AUROC for each DEG variant and dataset for data from the test set. Error bars indicate mean +/- s.e.m. across three random splits of the data. Note that human-human comparisons are not possible with AUROC, as humans produce binary labels, not probabilities. **(B)** AUROC for each DEG variant for individual behaviors from the Mouse-1 dataset. **(C-E)** Similar to (B), except for the Mouse-2 dataset, Mouse-3 dataset, Fly dataset. **(F)** DEG-medium performance measured as AUROC as a function of the percent of frames containing the behavior of interest in the training data. Each point is for one behavior for one split of the data, with three splits in total.

**Supplementary Figure 7:**
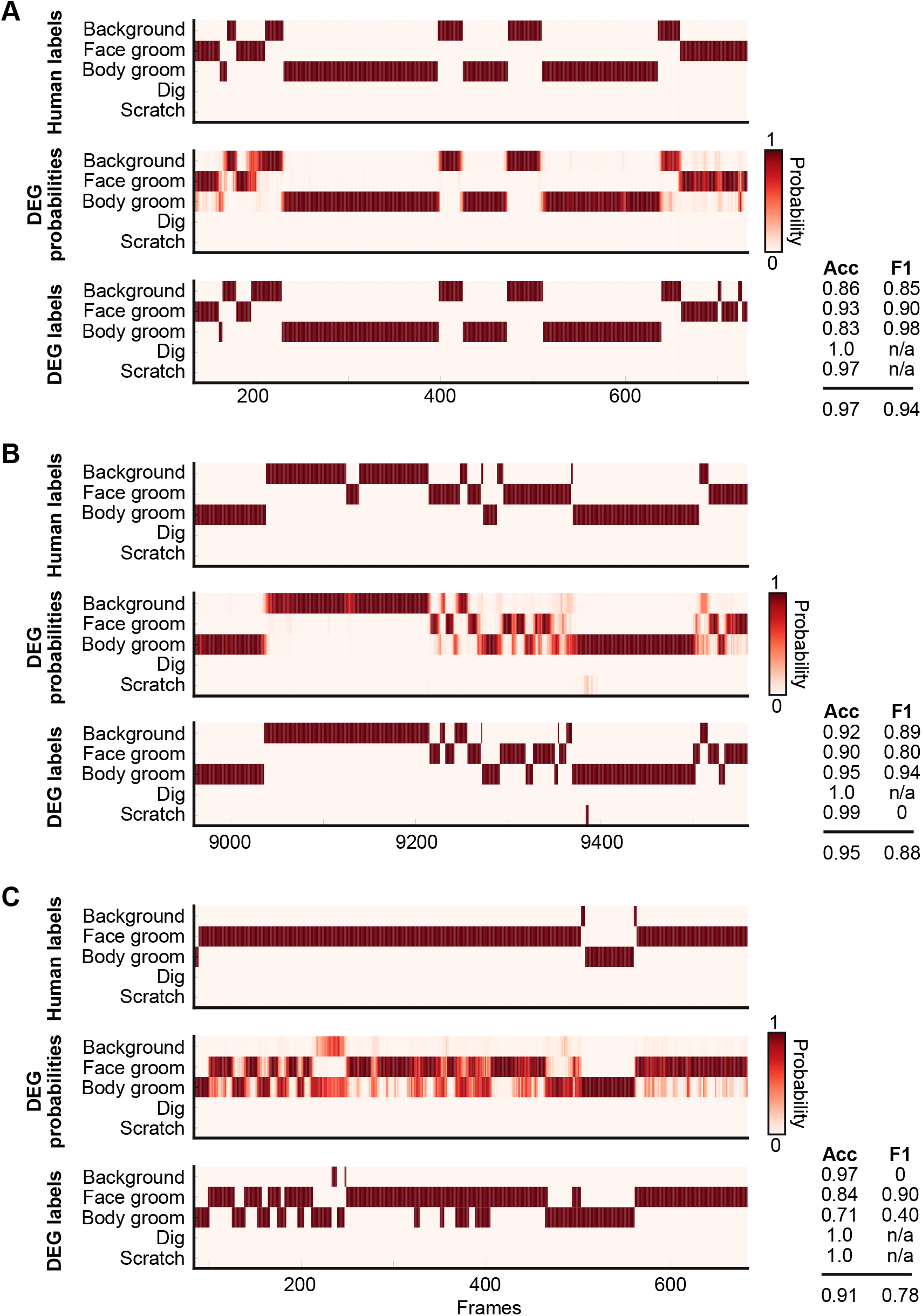
Ethogram examples for the Mouse-1 dataset. **(A)** An example ethogram with above-average performance, showing the human labels, estimated probabilities for each behavior from DEG-medium, and the thresholded and postprocessed predictions, for data from the test set. The accuracy and F1 score for each behavior are shown, along with the overall accuracy and overall F1 score. **(B-C)** Similar to (A), except for approximately average performance and below-average performance.

**Supplementary Figure 8:**
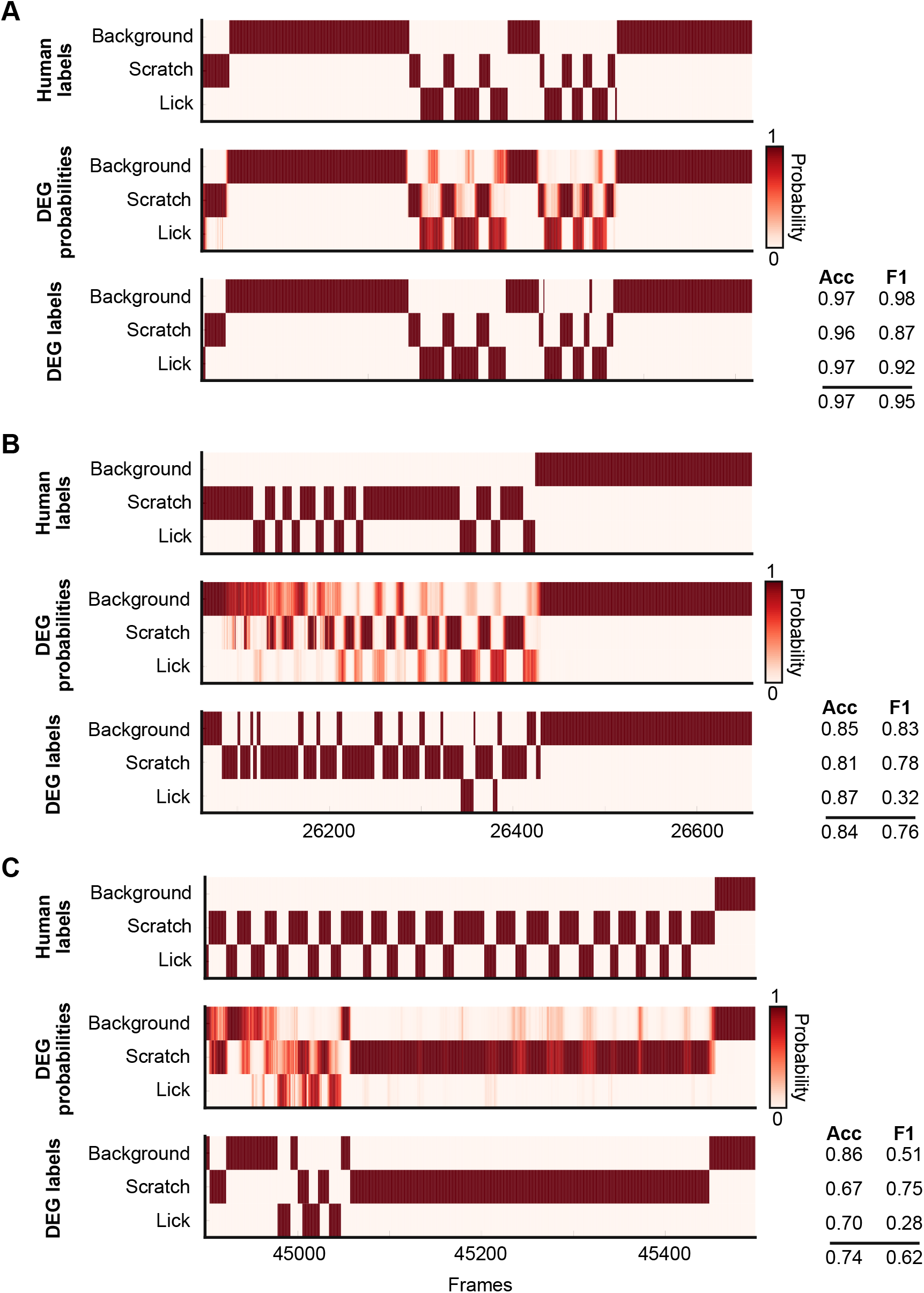
Ethogram examples for the Mouse-2 dataset. **(A)** An example ethogram with above-average performance, showing the human labels, estimated probabilities for each behavior from DEG-medium, and the thresholded and postprocessed predictions, for data from the test set. The accuracy and F1 score for each behavior are shown, along with the overall accuracy and overall F1 score. **(B-C)** Similar to (A), except for approximately average performance and below-average performance.

**Supplementary Figure 9:**
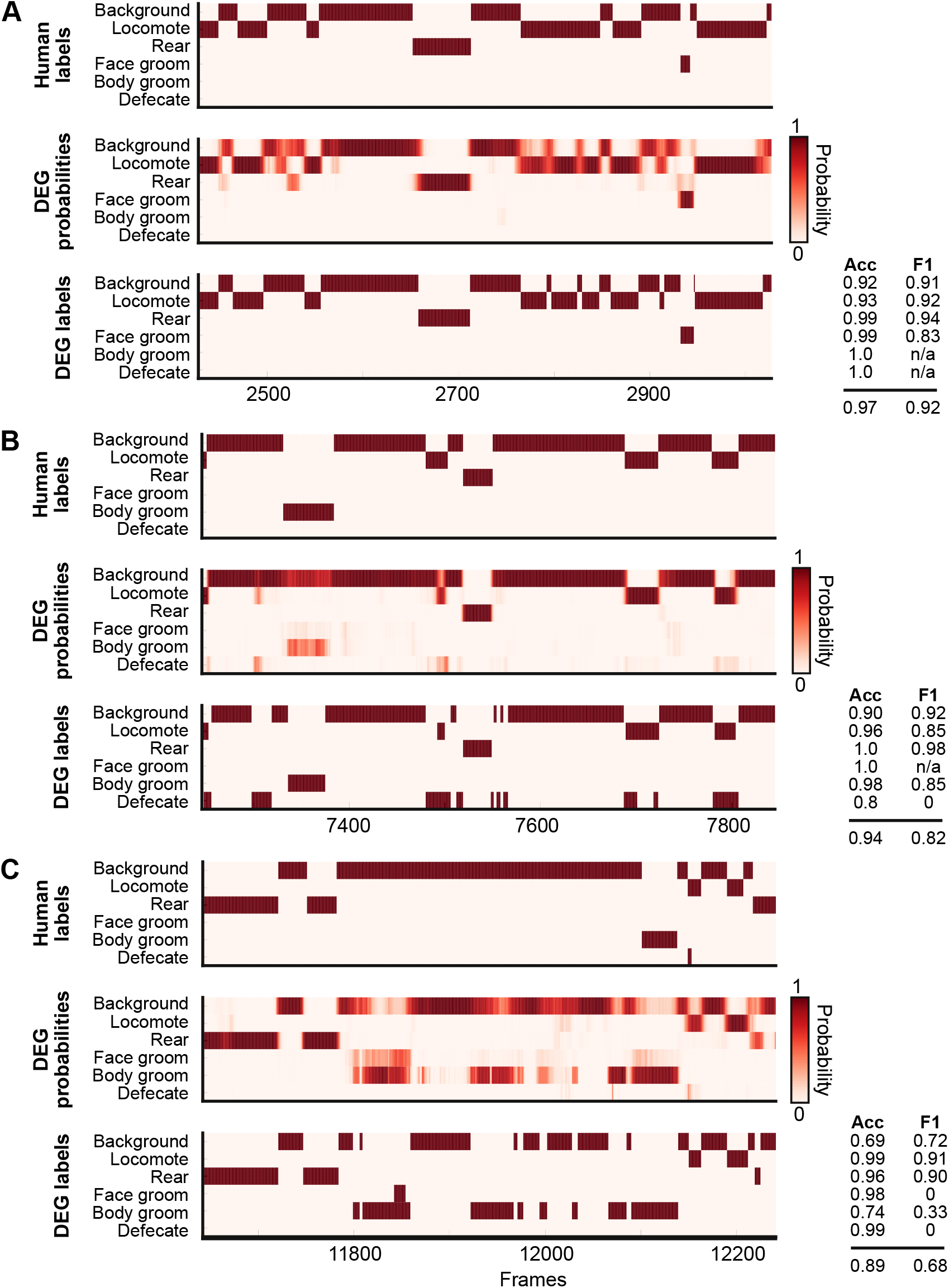
Ethogram examples for the Mouse-3 dataset. **(A)** An example ethogram with above-average performance, showing the human labels, estimated probabilities for each behavior from DEG-medium, and the thresholded and postprocessed predictions, for data from the test set. The accuracy and F1 score for each behavior are shown, along with the overall accuracy and overall F1 score. **(B-C)** Similar to (A), except for approximately average performance and below-average performance.

**Supplementary Figure 10:**
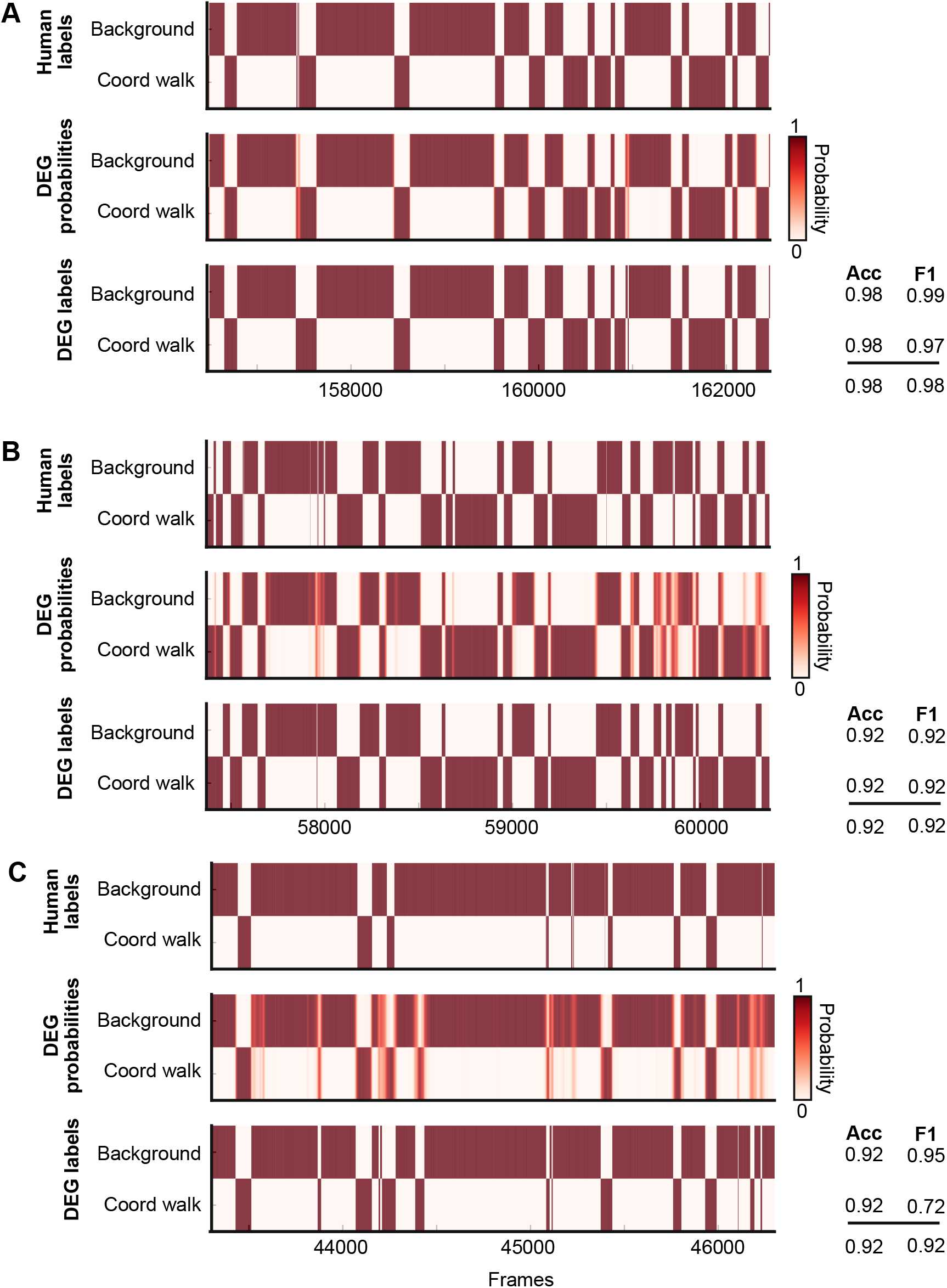
Ethogram examples for the Fly dataset. **(A)** An example ethogram with above-average performance, showing the human labels, estimated probabilities for each behavior from DEG-medium, and the thresholded and postprocessed predictions, for data from the test set. The accuracy and F1 score for each behavior are shown, along with the overall accuracy and overall F1 score. **(B-C)** Similar to (A), except for approximately average performance and below-average performance.

**Supplementary Figure 11:**
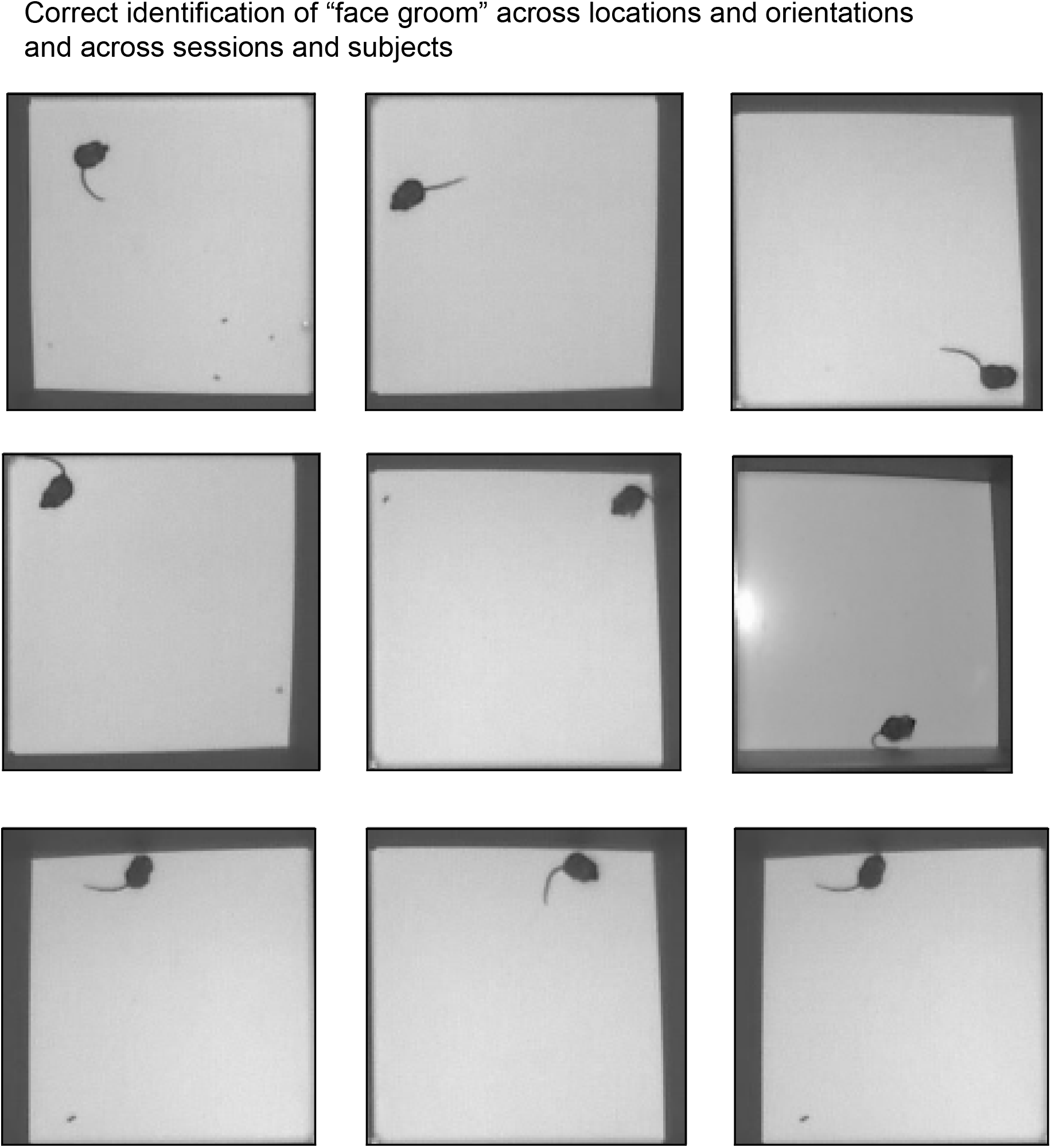
DeepEthogram exhibits position and heading invariance. Nine randomly selected examples of the “face groom” behavior from the Mouse-3 dataset. All examples were identified as “face groom” by DEG-medium. The examples include different videos and different mice.

## Notes

https://github.com/jbohnslav/deepethogram

### Competing Interest Statement

The authors have declared no competing interest.

